# ASHIC: Hierarchical Bayesian modeling of diploid chromatin contacts and structures

**DOI:** 10.1101/2020.08.29.273722

**Authors:** Tiantian Ye, Wenxiu Ma

## Abstract

The recently developed Hi-C technique has been widely applied to map genome-wide chromatin interactions. However, current methods for analyzing diploid Hi-C data cannot fully distinguish between homologous chromosomes. Consequently, the existing diploid Hi-C analyses are based on sparse and inaccurate allele-specific contact matrices, which might lead to inaccurate modeling of diploid genome architecture.

Here, we present ASHIC, a hierarchical Bayesian framework to model allele-specific chromatin organizations in diploid genomes. We developed two models under this Bayesian framework: the Poisson-multinomial (ASHIC-PM) model and the zero-inflated Poisson-multinomial (ASHIC-ZIPM) model. The proposed ASHIC methods impute allele-specific contact maps from diploid Hi-C data and simultaneously infer allelic 3D structures. Through simulation studies, we demonstrated that our methods outperformed existing approaches, especially under low coverage and low SNP density conditions. Additionally, we applied ASHIC-ZIPM to a published diploid mouse Hi-C data and studied the active/inactive X chromosomes and the *H19/Igf2* imprinting region. In both cases, our method produced fine-resolution diploid chromatin maps and 3D structures, and provided insights into the allelic chromatin organizations and functions. To summarize, our work provides a statistically rigorous framework for investigating fine-scale allele-specific chromatin conformations.

The ASHIC software is publicly available at https://github.com/wmalab/ASHIC.

## 1 INTRODUCTION

The three-dimensional (3D) organization of chromatin in the nucleus plays an essential role in gene regulation [1]. The recently developed chromosome conformation capture coupled with high-throughput sequencing (Hi-C) technique [2–4] and its variants [5–7] have been widely applied to map genome-wide chromatin interactions and to elucidate the principles of spatial genome architecture. The Hi-C experiment yields a genome-wide chromatin contact matrix; each entry (*i,j*) in the matrix represents the contact frequency between two loci *i* and *j* in the genome. The mapping and subsequent analyses of genome-wide Hi-C contact matrices in various organisms have demonstrated that the gene expression is tightly regulated by chromatin interactions at multiple scales ranging from active/inactive chromosomal compartments and sub-compartments [2, 6], to topologically associated domains (TADs) [8], and fine-scale chromatin loops [5, 6].

One hindrance of current Hi-C data analysis is the lack of allele-specific modeling for diploid genomes. Most mammalian genomes are diploid, in which the genome contains two sets of each chromosome—a maternal and a paternal copy. Hence, a chromatin contact observed between two genomic loci in the reference (haploid) genome may correspond to four distinct yet indistinguishable chromatin interactions in the diploid genome. For example, a chromatin contact mapped to a loci pair (*i, j*) on the same chromosome in the reference genome could be either an intra-chromosomal contact (*m_i_, m_j_*) on the maternal allele, or an intra-chromosomal contact (*p_i_, p_j_*) on the paternal allele, or inter-homologous contacts (*m_i_, p_j_*) or (*p_i_, m_j_*) (Figure 1A). However, the majority of existing Hi-C analyses on diploid genomes do not distinguish between homologous chromosomes. As a result, current analyses are based on an aggregated contact matrix generated with mixed signals of maternal and paternal chromatin contacts, which could result in the false identification of significant chromatin interactions and an inaccurate understanding of the diploid genome architecture. Therefore, statistical methods for rigorous and accurate modeling of diploid Hi-C data are needed to facilitate a detailed understanding of the mechanisms of chromatin organization and gene regulation.

**Figure 1:**
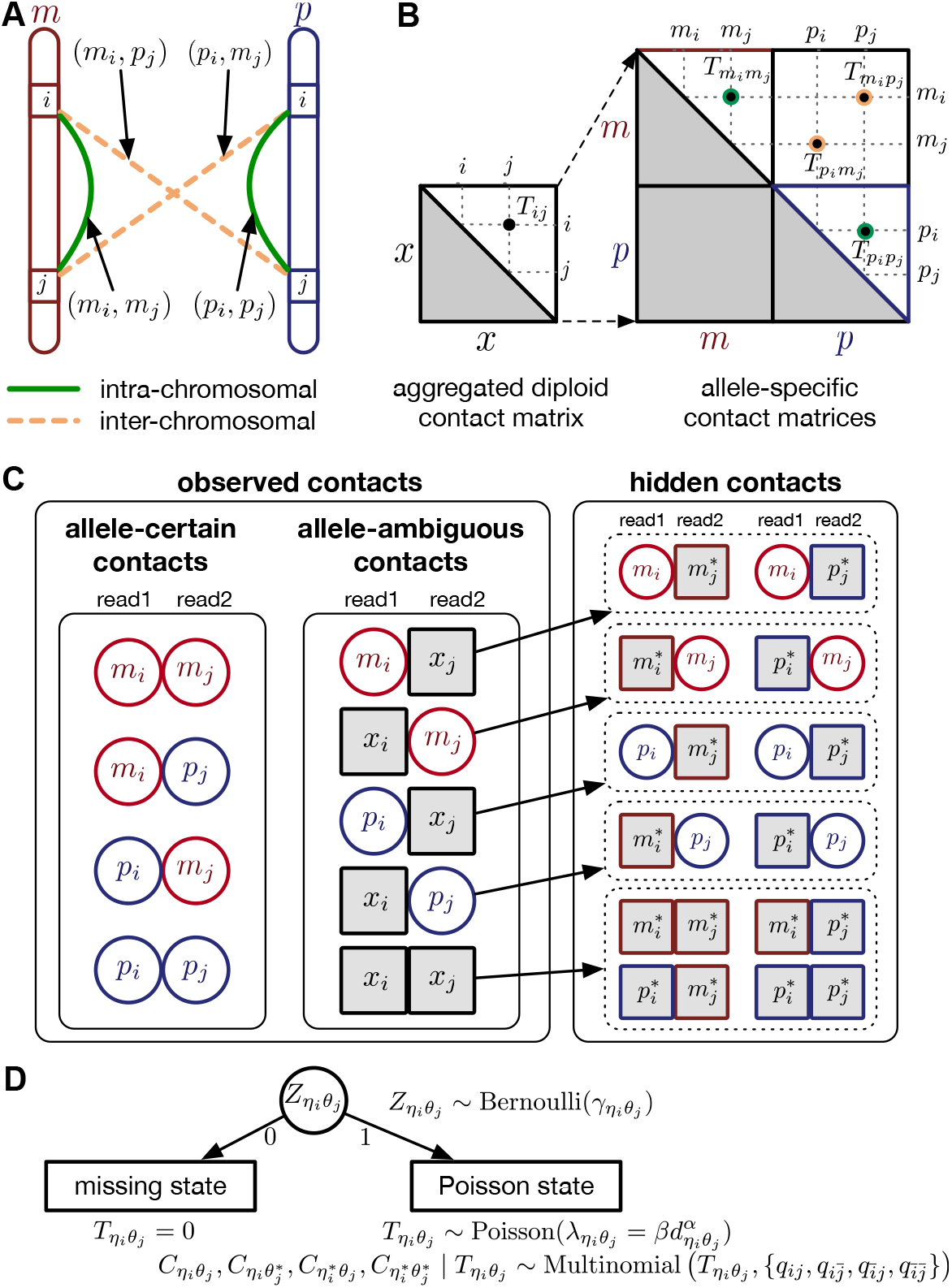
Overview of allele-specific modeling of diploid Hi-C data. **(A)** Diploid contact (*i, j*) is a combination of four distinct allele-specific contacts (*m_i_, m_j_*), (*m_i_, p_j_*), (*p_i_, m_j_*), and (*p_i_, p_j_*). **(B)** Reconstruction of allele-specific diploid contact matrix. **(C)** Observed allele-specific contacts between bins *i* and *j* can be decomposed into observed allele-certain contacts 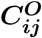, observed allele-ambiguous contacts 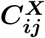. We aim to decompose 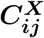 and infer the hidden contacts 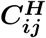, and impute the true allele-specific contacts **T_j_. (D)** Illustration of the hierarchical Bayesian ASHIC-ZIPM model.

Recently, several methods have been developed to obtain allele-specific chromatin contact matrices and/or allelic 3D structures from diploid Hi-C data [6, 9–14]. These methods use heterozygous single nucleotide polymorphisms (SNPs) to identify the allele identity of chromatin interactions. Specifically, a Hi-C contact is a mate-pair with two read ends representing the two interacting chromatin fragments. If a read end overlaps with SNPs for which the allele identity can be determined, we term it an allele-certain read. For example, a read containing maternal-specific SNP(s) is assigned to the maternal allele; similarly a read containing paternal-specific SNP(s) is assigned to the paternal allele. In addition, reads without SNPs are allele-ambiguous reads. Based on the allele identity of the paired ends, we can then categorize diploid Hi-C contacts into three groups: both-end allele-certain contacts, one-end allele-ambiguous contacts, and both-end allele-ambiguous contacts.

Without a statistically rigorous allele inference method, many previous studies applied either an “allele-certain” or a “mate-rescue” strategy to reconstruct the allele-specific contact maps in diploid genomes. First, in the allele-certain approach, only both-end allele-certain contacts are used [6, 11]. However, the both-end allele-certain contacts only account for a small portion of the total chromatin contacts (Supplementary Table S1). For example, in the patski (BL6×*Spretus*) cell line of which the SNP density is approximately 1 per 75 bp, the proportion of both-end allele-certain contacts in a typical Hi-C dataset is about 35.6%. Whereas, in the human GM12878 cell line of which the SNP density is approximately 1 per 1700 bp, the both-end allele-certain proportion drops to 0.14%. Consequently, the diploid contact matrices obtained by such an allele-certain approach is often sparse and of low resolution.

To overcome the low-coverage issue of the allele-certain approach, several diploid Hi-C studies adopted a straightforward mate-rescue strategy to infer the allele identity of one-end allele-ambiguous contacts, i.e., the allele-ambiguous end of such contact is assigned to the same allele as its mate-end [10, 12, 15]. This mate-rescue method attempts to recover one-end allele-ambiguous contacts, which varies approximately from 5.7% (in the case of GM12878 cells) to 43.3% (in the case of patski cells) of the total contacts (Supplementary Table S1). However, one-end allele-ambiguous contacts are all assumed to be intra-chromosomal contacts in the results of the mate-rescue approach. Such false assumption would lead to inaccurate contact maps, especially in the regions where inter-chromosomal interactions are observed across chromosomal territories.

Since the mate-rescue method fails to infer inter-chromosomal interactions from one-end allele-ambiguous contacts, Tan et al. [13] proposed an iterative two-stage imputation algorithm Dip-C for modeling single-cell diploid Hi-C data. In the first imputation stage, one-end allele-ambiguous contacts are phased using an *ad hoc* voting procedure by their neighborhood on the contact matrix. In the second imputation stage, the assignment of allele-ambiguous contacts is refined by the 3D structures. The Dip-C method can be viewed as an advanced mate-rescue method, as it leverages additional information from both contact matrices and 3D structures to infer allele-ambiguous contacts. However, the Dip-C method is specifically designed for single-cell Hi-C data therefore may not adapt well to bulk Hi-C data. Moreover, it uses a deterministic voting strategy to assign allele-ambiguous contacts, which does not provide a probabilistic model of all possible allele origins.

One common drawback of the allele-certain and mate-rescue methods is that they do not utilize both-end allele-ambiguous contacts, which represent a substantial proportion in the total diploid contacts, ranging from 21.1% (patski) to 94.1% (GM12878) (Supplementary Table S1). Inferring the allele identity of both-end allele-ambiguous contacts remains a significant challenge. To date, few methods have been developed to address this problem. The Dip-C method [13] attempts to impute only inter-chromosomal rather than intra-chromosomal both-end allele-ambiguous contacts. Thus, it does not produce a fully recovered diploid contact map. In addition, our previously proposed Poisson-Gamma model [9] imputes both one-end and both-end allele-ambiguous contacts, and estimates the diploid contact matrices by an iterative expectation-maximization (EM) algorithm [16]. The Poisson-Gamma method takes advantage of all diploid contacts. However, it does not predict diploid 3D structures nor use the structures to assist the inference of allele-ambiguous contacts. As a result, the Poisson-Gamma model might not work robustly in fine-resolution analyses. Furthermore, Cauer et al. [14] developed diploid-PASTIS, an extension of the PASTIS model [17], to infer the diploid chromatin structures. Diploid Hi-C contacts are modeled as Poisson variables, and the optimal diploid structures are solved by maximizing the likelihood function with additional structural constraints. The diploid-PASTIS method is specifically designed to model diploid 3D structures, but does not infer the allele-ambiguous contacts to impute the full diploid contact matrices.

To tackle the aforementioned challenges, we developed a hierarchical Bayesian framework for Allele-Specific diploid Hi-C modeling, named ASHIC. Briefly, allele-specific contact counts are modeled as Poisson-multinomial random variables (referred as the ASHIC-PM model) and diploid contact matrices and 3D structures are estimated via an EM algorithm. Moreover, to overcome the sparsity issue of diploid Hi-C contact maps, we proposed a zero-inflated version of the ASHIC-PM method, namely the zero-inflated Poisson-multinomial model (in short, ASHIC-ZIPM). Both ASHIC models can completely dissect all diploid Hi-C contacts into allele-specific contact maps, while simultaneously reconstruct 3D homologous chromosomal structures. To the best of our knowledge, our ASHIC methods are the first methods that fully impute all allele-ambiguous contacts and infer both the diploid contact matrices and allelic 3D structures.

We thoroughly evaluated our methods through a series of simulation studies and demonstrated that our ASHIC methods outperformed existing allele-certain and mate-rescue approaches under settings with varied sequencing coverage, SNP density, and homologous structural similarity. We also applied the ASHIC-ZIPM method to a published diploid mouse Hi-C data [15]. We successfully confirmed that our predicted diploid contact maps and 3D structures of the homologous X chromosomes exhibited distinct chromatin conformations, where the inactive X demonstrated the bipartite superdomains [9]. Furthermore, we studied fine-scale chromatin organizations of the imprinted *H19/Igf2* region at 10 kb resolution and revealed distinct parental-specific chromatin interactions anchored at *H19* and *Igf2*. With the fully imputed diploid contact matrices, we un-covered a maternal-specific sub-TAD organization at the *H19/Igf2* imprinting region.

## 2 MATERIALS AND METHODS

### 2.1 Notations of allele-specific chromatin contacts in diploid genomes

Let *m* and *p* denote a homologous chromosomal pair with same length *n* in a diploid genome. To construct the diploid Hi-C contact frequency matrix, we partition the chromosomes into fixed-size non-overlapping bins and count chromatin contacts observed between each bin pair. In the diploid setting, chromatin contacts between the bins *i* and *j* can result from four distinct events: (*m_i_, m_j_*), (*p_i_, p_j_*), (*m_i_, p_j_*), or (*p_i_, m_j_*), where (*m_i_, m_j_*) and (*p_i_, p_j_*) are intra-chromosomal contacts on chromosome *m* and *p*, respectively, and (*m_i_, p_j_*) and (*p_i_, m_j_*) are inter-chromosomal contacts between the homologous chromosomes (Figure 1A). Therefore, the aggregated contact frequency *T_i,j_* between the bins *i* and *j* can be calculated as follows: *T_ij_* ∑_*η*_ ∑_*θ*_ *T_η_i_θ_j__*, where *T_η_i_θ_j__* is the unknown true allele-specific contact frequency between *η_i_* (bin *i* on chromosome *η*) and *θ_j_* (bin *j* on chromosome *θ*) that we aim to estimate, *η, θ* ∈ {*m, p*}, 1 ≤ *i, j* ≤ *n* (Figure 1B).

Using heterozygous SNPs, we can classify single-end reads into three categories: reads containing allele-*m*-specific SNPs, reads containing allele-*p*-specific SNPs, and reads containing no SNPs. We refer to the first two categories as allele-certain reads while the last category as allele-ambiguous reads. Furthermore, since Hi-C contacts are paired-end reads, each end of the mated pair can either be allele-certain or allele-ambiguous. Let *C_η_i_θ_j__* indicate the frequency of both-end allele-certain contacts between the bins ηi and *θ_j_*. In addition, we specify 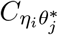. to be the contact frequency between *η_i_* and *θ_j_* where the allele identity of *η_i_* is known but the allele identity of *θ_j_* is unknown. In other words, one end of the Hi-C contact is from *θ_j_*; however, the read does not overlap with any SNPs. Therefore the allele identity of *θ_j_* remains unknown. Similarly, we use 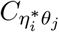 when the allele identity of *η_i_* is unknown and 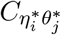 when the allele identity of both ends are unknown. Hence, the true allele-specific contact frequency *T_η_i_θ_j__*. equals to the sum of the following four components:

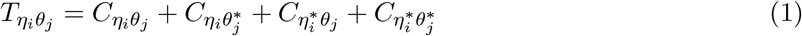

In diploid Hi-C data, we cannot directly observe 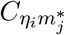 and 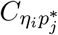 since the read end mapped to the bin *j* is allele-ambiguous and hence, it cannot be distinguished between *m_j_* and *p_j_*. As a result, the observed Hi-C contacts contain the following types of allele-ambiguous contacts:

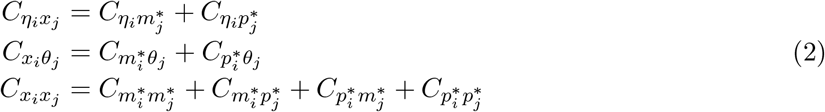

where *x* indicates that the allele identity of a read end is unknown. We refer to *C_η_i_x_j__* and *C_x_i_θ_j__* as one-end allele-ambiguous contacts and *C_x_i_x_j__* as both-end allele-ambiguous contacts (Figure 1C).

In summary, we define ***C^O^*** = {*C_η_i_θ_j__*} as the observed allele-specific contact frequencies, ***C^X^*** = {*C_η_i_x_j__, C_x_i_θ_j__, C_x_i_x_j__*} as the observed allele-ambiguous contact frequencies (LHS in eq. (2)), and 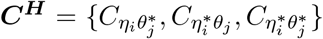 as the unobserved (hidden) allele-specific contact frequencies (RHS in eq. (2)). Our goal is to decompose **C^X^** and infer **C^H^** in order to impute the true allele-specific frequencies ***T*** = {*T_η_i_θ_j__*} by eq. (1) (Figure 1C).

### 2.2 Hierarchical Bayesian modeling of diploid chromatin contact maps and 3D structures

To model diploid Hi-C data, we propose a hierarchical Bayesian modeling framework for imputing the allele-specific chromatin contacts and reconstructing the allelic 3D structures. Specifically, we model the generation of allele-specific contacts with either a Poisson-multinomial process (the ASHIC-PM model) or a zero-inflated Poisson-multinomial process (the ASHIC-ZIPM model) for the inference of diploid contact matrices and 3D structures. The ASHIC-ZIPM model is a zero-inflated version of ASHIC-PM and it explicitly accounts for the excessive zeros observed in Hi-C contact matrices.

Here, we describe ASHIC-ZIPM, the zero-inflated version of our hierarchical Bayesian model and the corresponding EM algorithm for model fitting. The details for the ASHIC-PM model are available in Supplementary Methods.

#### 2.2.1 Modeling true allele-specific contact frequencies from diploid 3D structures

We adopt the coarse-grained polymer model [18] to represent the chromosomal structures. Each bin in the genome is represented as a bead in the 3D space, and each chromosome can be viewed as a chain of beads. Specifically, we denote ***X_m_*** and ***X_p_*** to be the 3D coordinates of the homologous chromosomes *m* and *p*, respectively, where ***X_m_, X_p_*** ∈ *R*^3×*n*^. Let ***x_η_i__*** and ***x_θ_j__*** to be the 3D coordinates of beads *η_i_* and *θ_j_*, respectively, where *η, θ* ∈ {*m, p*}. According to polymer physics [17, 19], the contact frequency *T_η_i_θ_j__* between *η_i_* and *θ_j_* is inversely correlated with their spatial distance *d_η_i_θ_j__*, following a power-law decay function. That is, 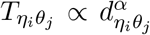, where *α* < 0 is the exponent of the distance-decay function, and *d_η_i_θ_j__* = ‖***x****_η_i__ − x_θ_j__*‖_2_ is the Euclidean distance between beads *η_i_* and *θ_j_*.

Similar to the PASTIS method [17], we model the true allele-specific contact frequency *T_η_i_θ_j__* as a Poisson random variable, *T_η_i_θ_j__* ~ Poisson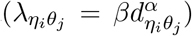, where *β* is a scaling factor (the ASHIC-PM model, Supplementary Methods). Furthermore, to account for the excessive zeros in Hi-C contact matrices, we propose to use a zero-inflated Poisson (ZIP) distribution to model the contact counts (the ASHIC-ZIPM model) (Figure 1D).

In the ASHIC-ZIPM model, we assume that *T_η_i_θ_j__* follows a ZIP distribution.

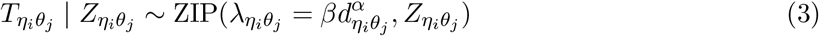

Different from the ASHIC-PM model, here we introduce *Z_η_i_θ_j__*, a latent binary variable to indicate whether *T_η_i_θ_j__* is generated from the Poisson state (*Z_η_i_θ_j__* = 1, *T_η_i_θ_j__* ~ Poisson(λ_*η_i_θ_j_*_)) or the missing state (*Z_η_i_θ_j__* = 0, *T_η_i_θ_j__* = 0).

Furthermore, we assume that *Z_η_i_θ_j__* follows a Bernoulli prior with a success probability *γ_η_i_θ_j__*.

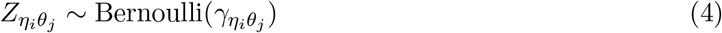

For intra-chromosomal contacts (where *η = θ*), *γ_η_i_θ_j__* is a function of the corresponding genomic distance. For inter-chromosomal contacts (η ≠ θ), *γ_η_i_θ_j__* is set to a constant.

In other words, the true allele-specific contact frequency T*η_i_θ_j_* is a mixture of two states. In the Poisson state (with probability *γ_η_i_θ_j__*), *T_η_i_θ_j__* follows a Poisson distribution; whereas in the missing state (with probability 1 − *γ_η_i_θ_j__*), *T_η_i_θ_j__* = 0. The *γ_η_i_θ_j__* parameter acts as a weight between the Poisson and missing states. Note that when all latent variables ***Z*** = {*Z_η_i_θ_j__*} are equal to 1, the ASHIC-ZIPM model reduces to the ASHIC-PM model. Hence, ASHIC-PM is a special case of ASHIC-ZIPM.

#### 2.2.2 Modeling allele-identifiable probability and allele-ambiguous contact counts

As discussed above, we cannot directly observe the allele identity of all diploid Hi-C contacts. We use *q_i_* to denote the allele-identifiable probability of bin *i* in the genome, i.e., if a single-end read is mapped to bin *i*, the probability that the read overlaps with SNP(s) (and therefore can be distinguished between alleles *m* and *p*) is *q_i_*. Consequently, assuming that bins *i* and *j* are independent, the probabilities that a paired-end contact between the bins *i* and *j* is both-ends allele-certain (*q_ij_*), one-end allele-ambiguous at bin 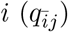, one-end allele-ambiguous at bin 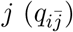, and both-end allele-ambiguous 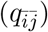 can be calculated as follows:

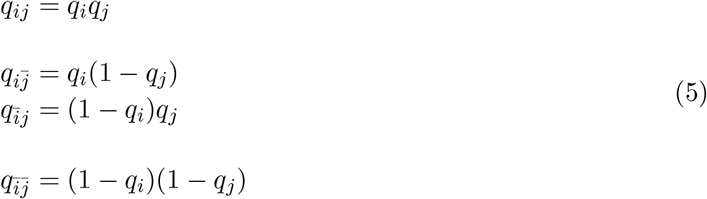

Recall in eq. (1), the true allele-specific contact frequency *T_η_i_θ_j__* can be expressed as the sum of one observed allele-certain contact frequency *C_η_i_θ_j__* and three unobserved hidden allele-specific contact frequencies 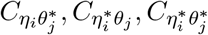. We assume that the decomposed allele-specific contact frequencies follow a multinomial distribution.

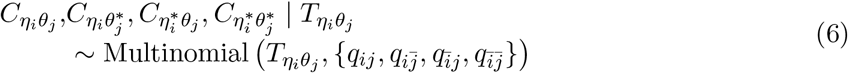

Based on the above assumptions, we derive that 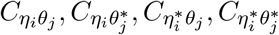 given *Z_η_i_θ_j__* are mutually conditional independent ZIP random variables. Consequently, we demonstrate that the observed allele-ambiguous contact frequencies *C_η_i_x_j__, C_x_i_θ_j__*, and *C_x_i_x_j__* are ZIP random variables and mutually conditional independent given ***Z*** (Supplementary Methods).

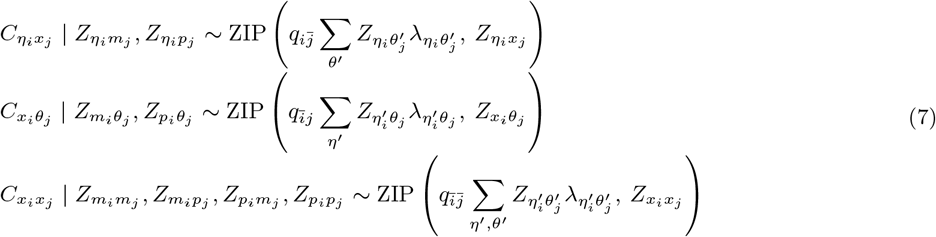

where *Z_η_i_χ_j__* := *Z_η_i_m_j__* or *Z_η_i_p_j__*, i.e., *Z_η_i_χ_j__* = 1 if *Z_η_i_m_j__* = 1 or *Z_η_i_p_j__* = 1. Similarly, we denote *Z_χ_i_θ_j__* := *Z_m_i_θ_j__* or *Z_p_i_θ_j__, Z_x_i_x_j__* := *Z_m_i_m_j__* or *Z_m_i_p_j__* or *Z_p_i_m_j__* or *Z_p_i_p_j__*.

#### 2.2.3 Incorporating bias factors

Real Hi-C data contains various types of systematic biases. Similar to the ICE method [20], we assume that the bias of observing contacts between bins *η_i_* and *θ_j_* can be factorized as the product of the bias factors *b_η_i__* and *b_θ_j__* of the two bins, respectively. Hence, we can incorporate bias factors into the ASHIC-ZIPM model as follows:

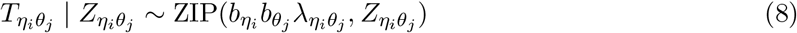

Our ASHIC software provides an option to estimate the bias factors from real diploid Hi-C data and to incorporate them into our model (Supplementary Methods).

### 2.3. Estimating allele-specific chromatin structures and contact frequencies via EM algorithm

We design an EM algorithm to simultaneously infer 3D structures and estimate model parameters. The EM algorithm for the ASHIC-ZIPM model is briefly outlined below. Details of the EM algorithms for both ASHIC-PM and ASHIC-ZIPM are available in Supplementary Methods.

In the ASHIC-ZIPM model, the parameter space contains the homologous chromosome structures ***X_m_*** ∈ *R*^3×*n*^ and ***X_p_*** ∈ *R*^3×*n*^, the distance-decay exponent *α*, the scaling factor *β*, the hyper parameter ***γ*** = {*γ_η_i_θ_j__*} (for the Bernoulli prior distribution of the Poisson state latent variables ***Z*** = {*Z_η_i_θ_j__*}), and the allele-identifiable probabilities ***q*** = {*q_k_*}, 1 ≤ *k ≤ n*. Note that in eq. (3), the ZIP parameter λ_*η_i_θ_j_*_ is a function of *α, β, **X**_m_* and ***X**_p_*. Here, we fix *α* and *β* in order to obtain a unique solution for ***X**_m_* and ***X**_p_*. Specifically, we use true estimate of *α* in simulations and set *α* = −3 in real data. We set *β* = 1 in both cases.

From the diploid Hi-C data, we can directly observe the allele-certain contacts ***C^O^*** and the allele-ambiguous contacts ***C^X^***. The unobserved latent variables include the hidden allele-specific contacts ***C^H^*** and the Poisson state latent variables ***Z***. The goal of the EM algorithm is to find the maximum likelihood estimate (MLE) of the model parameters, reconstruct the allele-specific 3D structures ***X_m_*** and ***X_p_***, and impute the true allele-specific contacts ***T***.

The complete likelihood of the observed data {***C^O^, C^X^***} and the unobserved latent variables {***C^H^, Z***} is

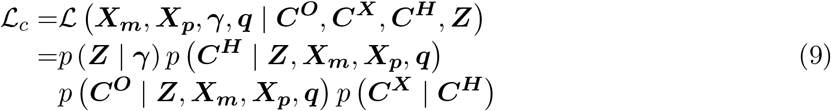

To solve the MLE of the marginal likelihood of observed data {***C^O^, C^X^***}, we propose an EM algorithm which applies the following two steps iteratively:

- Expectation step (E-step):

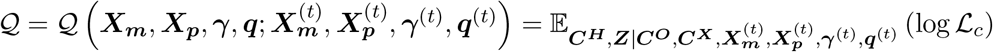
- Maximization step (M-step):

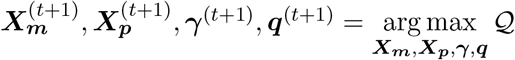

The pseudocode and detailed steps of the EM algorithm are available in Supplementary Methods. In particular, while estimating the homologous structures ***X**_m_* and ***X**_p_*, we develop an inter-homologous optimization strategy. Briefly, we first estimate ***X**_m_* and ***X**_p_* separately, then estimate the relative position between the two homologs to improve the final structure prediction (Supplementary Methods).

### 2.4. Simulation settings

#### 2.4.1 Simulating allele-specific chromatin contacts from homologous X chromosome structures

First, We considered the homologous X chromosomes as the ground truth and simulated diploid Hi-C datasets as described below. We assumed that the allele-specific chromatin contact frequencies follow the ASHIC-ZIPM model. The true model parameters *α_m_, α_p_, β, **γ***, and ***q*** were estimated from two published datasets on human GM12878 cells: (1) the predicted X chromosome structures from single-cell Hi-C data by Tan et al. [13], and (2) the allele-specific contact matrices from *in situ* bulk Hi-C data by Rao et al. [6], both at 100 kb resolution (Supplementary Methods, Supplementary Table S2).

At the default setting, we generated 10 simulated allele-specific Hi-C datasets with the scale factor 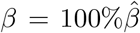 and the average allele-identifiable probability 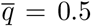. Subsequently, we kept other parameters fixed and generated 10 additional datasets for each of the decreased *β* values (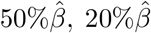, and 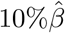), and another 10 datasets for each of the decreased 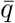 values (0.25, 0.1, and 0.05). In total, 70 diploid Hi-C datasets were generated in this simulation study. For each simulated dataset, we ran 10 random initializations and chose the result with the highest observed log-likelihood for performance evaluation and subsequent analyses.

#### 2.4.2 Simulating allele-specific chromatin contacts from identical chromosome structures

To study the effect of structural differences on the performance of our methods, we deployed a challenging simulation setting where we simulated diploid Hi-C datasets using two identical chromosome structures. Briefly, we duplicated the paternal (inactive) X chromosome structure ***X**_p_* predicted by Tan et al. [13] as the pseudo-maternal structure. Then we used the two identical chromatin structures as the ground truth and simulated diploid Hi-C datasets in a similar manner as previously described.

The relative position of these two identical structures was determined by a reversed structural superposition procedure. Using the original homologous structures ***X**_m_* and ***X**_p_*, we calculated the optimal translation vector ***υ*** and rotation matrix ***R*** using the Kabsch algorithm [21], such that the root-mean-square deviation (RMSD) between 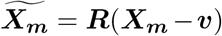 and ***X**_p_* was minimized. Then we duplicated ***X**_p_* and reversed the superposition of ***X**_p_* by 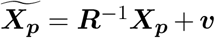. The resulting identical structures 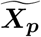 and ***X**_p_* was served as the pseudo-homologous chromosome structures in which the relative position between 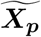 and ***X**_p_* remained approximately the same as the original homologous pair ***X**_m_* and ***X**_p_*.

### 2.5 Real Hi-C data processing and analysis

The published datasets used in our study are summarized in Supplementary Table S2. In particular, for diploid Hi-C data analysis, we used a wild-type patski dataset published by Bonora et al. [15] and focused on two chromosomal regions. The first one is the entire X chromosomes; the second one is the *H19/Igf2* imprinting region on chromosomes 7. For each region, we ran 20 random initializations with the ASHIC-ZIPM model and chose the one with the highest likelihood for subsequent analyses.

### 2.6 Convergence and running time

The convergence of the EM algorithm is defined as the relative increase of log-likelihood between two consecutive iterations is less than 10^−4^. We tested our ASHIC software using a single core on an Intel E5-2683v4 processor with 8GB memory allocation. In a typical simulation setting, two X chromosomes were partitioned into 3000+ bins at 100 kb resolution. With the default sequencing coverage 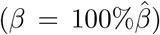 and SNP density 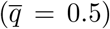 setting, both ASHIC-ZIPM and ASHIC-PM converged within 20 iterations (2 h). Lower coverage or lower SNP density requires more iterations. For example, when 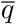 reduced to 0.05, the EM algorithm of ASHIC-ZIPM took about 50 iterations (8 h) to converge. When *β* decreased to 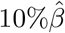, the EM algorithm of ASHIC-ZIPM underwent about 90 iterations (20 h) to converge.

### 2.7 Evaluation metrics

We used the following evaluation metrics in the simulation studies: the recovery rate (RR) for measuring the proportion of diploid Hi-C contacts recovered by each method, the imputation error rate (IER) and the stratum adjusted correlation coefficient (SCC) [22] for measuring the accuracy of imputed diploid contact matrices, the distance error rate (DER) and homologous distance error rate (HDER) for measuring the accuracy of predicted allelic 3D structures. Additionally, we calculated the recall, precision and *F*_1_ score to evaluate the allele-specific chromatin interactions identified from imputed diploid contact matrices. In the real data analysis of mouse X chromosomes, we used the bipartite index (BI) [9] to measure the bipartition organization of the inactive X chromosome, and calculated the radius of gyration (*R_g_*) to measure the compactness of both X chromosomes. The detailed definitions of these evaluation metrics can be found in Supplementary Methods.

## 3 RESULTS

### 3.1 Simulation studies on homologous X chromosomes

#### 3.1.1 Default simulation setting

We first evaluated the performance of the proposed ASHIC methods on simulated diploid Hi-C datasets of the homologous X chromosomes in human GM12878 cells. Of the two X chromosomes, the active X chromosome (denoted as Xa) is the maternal copy and the inactive X chromosome (denoted as Xi) is the paternal copy. We considered the 3D structures of Xa and Xi published by Tan et al. [13] as the ground truth and generated 10 simulated diploid Hi-C datasets at 100-kb resolution (see Methods). Each simulated dataset contained two intra-chromosomal contact matrices, one for Xa and one for Xi, as well as one inter-chromosomal contact matrix between Xa and Xi.

We compared our ASHIC-ZIPM and ASHIC-PM methods with two commonly used approaches for analyzing diploid Hi-C data. The first approach is the allele-certain method that uses only both-end allele-certain contacts [6, 11]. The second approach is the mate-rescue method that combines both-end allele-certain contacts with one-end allele-ambiguous contacts by assigning the allele-ambiguous read-end to the same allele as the allele-certain mate-end [10, 12, 15].

To evaluate the imputation of diploid Hi-C contact maps, we first calculated the proportion of allele-specific contacts recovered by each method (Supplementary Table S3). At the default sequencing coverage 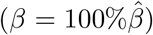 and SNP density 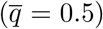 setting, the allele-certain and mate-rescue approaches recovered evidently smaller proportion of diploid chromatin contacts (25.65% and 75.55%, respectively) compared to the ASHIC-ZIPM and ASHIC-PM methods that were able to recover all one-end and both-end allele-ambiguous reads, thereby achieving 100% full recovery rate.

Next, we sought to assess the accuracy of the imputed allele-specific contact matrices. Recent studies have demonstrated that the genomic distance dependence and sequencing depth have confounding effects on measuring the similarity between Hi-C contact matrices [22]. To account for these confounding factors, we computed the stratum adjusted correlation coefficient (SCC) using the `HiCRep` package [22] to measure the similarity between the imputed contact matrices and true matrices (Figure 2A). We observed that the imputed diploid matrices obtained by ASHIC-ZIPM and ASHIC-PM had near-perfect SCC values of 0.9997 and 0.9996, respectively; whereas mate-rescue and allele-certain methods demonstrated lower SCC values of 0.9733 and 0.8100, respectively. ASHIC-ZIPM showed a significantly higher SCC values than ASHIC-PM (*p*-value = 2.53 × 10^−3^, one-sided paired Wilcoxon signed-rank test). In addition, ASHIC-ZIPM performed significantly better than the allele-certain and mate-rescue methods (*p*-values = 2.53 × 10^−3^, one-sided paired Wilcoxon signed-rank tests). Note that *p* = 2.53 × 10^−3^ is the smallest possible *p*-value given the sample size.

**Figure 2.**
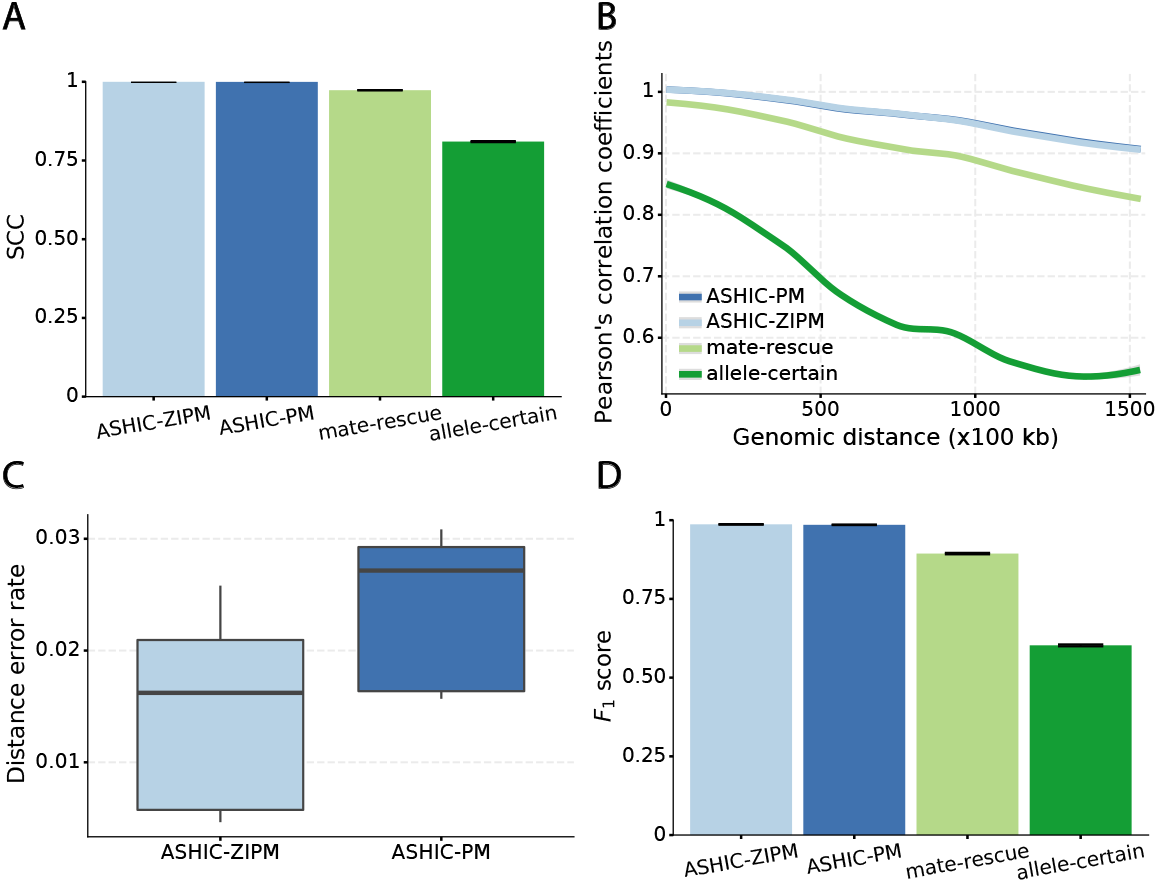
Evaluation on simulated homologous X chromosome (Xa/Xi) data. **(A)** Stratum-adjusted correlation coefficients (SCCs) and **(B)** and Pearson’s correlation coefficients (PCCs) between the imputed diploid contact matrices and the true contact matrices. The PCC curves are smoothened using the locally weighted LOESS method. **(C)** Distance error rates between the predicted allelic 3D structures and the true structures. **(D)** *F*_1_ scores of the identified allele-specific chromatin interactions.

The SCC statistic is a weighted average of the Pearson’s correlation coefficients (PCCs) across different genomic distances [22]. To breakdown the effect of genomic distance, we computed the PCCs between the imputed contact matrices and the true matrices at different genomic distances (Figure 2B). As expected, the PCC values decreased as the genomic distance increased for all four methods. We observed that the ASHIC-ZIPM and ASHIC-PM methods demonstrated similar PCC values across all genomic distances. In addition, the ASHIC-ZIPM and ASHIC-PM methods outperformed the allele-certain and mate-rescue approaches by large margin, especially at large genomic distances. Taken together, the SCC and PCC results showed that our ASHIC methods can accurately impute allele-specific contact matrices. Moreover, the imputation accuracy outperformed the allele-certain and mate-rescue approaches, especially for long-range contacts.

In addition to imputing diploid Hi-C contact matrices, the ASHIC-ZIPM and ASHIC-PM methods also predict allele-specific 3D structures. To evaluate the accuracy of the predicted allelic structures, we calculated the distance error rates between the predicted structures and the ground truth (Figure 2C). We observed that ASHIC-ZIPM yielded significantly lower distance error rates and thereby, more accurate allelic 3D structures than those obtained by ASHIC-PM (*p*-value = 2.53 × 10^−3^, one-sided paired Wilcoxon signed-rank test).

Furthermore, we investigated whether the imputed diploid contact matrices can facilitate the detection of allele-specific chromatin interactions. First, we called significant interactions using the Fit-Hi-C package [23] on the true diploid contact matrices. We subsequently defined the maternal-specific interactions as the interactions that were called only from the true maternal matrix but not from the paternal matrix. The paternal-specific interactions were defined accordingly. The final set of true allele-specific interactions was defined as the union of both monoallelic sets, which contained 9061.5 interactions on average (Supplementary Table S5). Following the same procedure, we then identified the allele-specific interactions from the imputed diploid contact matrices resulting from the four methods, separately. We evaluated the identified allele-specific interactions from each method using three metrics: precision, recall, and their harmonic mean *F*_1_ score (Figure 2D, Supplementary Figure S1). ASHIC-ZIPM and ASHIC-PM maintained the highest *F*_1_ scores of 0.9867 and 0.9853, respectively. In addition, ASHIC-ZIPM significantly outperformed mate-rescue (*F*_1_ = 0.8940) and allele-certain (*F*_1_ = 0.6024) in terms of the *F*_1_ scores (*p*-values = 2.53 × 10^−3^, onesided paired Wilcoxon signed-rank tests). The low *F*_1_ scores of the mate-rescue and allele-certain methods were primarily contributed by their low recall rates (Supplementary Figure S1), which was a result of their low recovery rates of allele-ambiguous contacts (Supplementary Table S3).

Collectively, our comparisons have demonstrated that the proposed ASHIC-ZIPM and ASHIC-PM methods outperformed the existing mate-rescue and allele-certain approaches with respect to the recovery rate of allele-ambiguous contacts, the accuracy of imputed diploid contact matrices and predicted allelic 3D structures, and the ability to facilitate the detection of allele-specific chromatin loops. In addition, ASHIC-ZIPM demonstrated a better performance overall than that of ASHIC-PM, especially in the prediction of allelic 3D structures. To further evaluate the performance of these methods under different circumstances, we conducted a series of additional simulation experiments by adjusting three major factors: sequencing coverage, SNP density, and homologous structural similarity.

#### 3.1.2 Performance on low sequencing coverage data

The sequencing coverage of Hi-C contact matrices is a major factor that can affect the performance of the diploid Hi-C methods. An observed zero entry in the Hi-C contact matrix can be either a “true” zero as a result of no physical contact between the pair of chromatin fragments, or a “missing” zero as a result of insufficient sequencing coverage. Lower sequencing depth of Hi-C experiments yields lower-coverage and sparse contact matrices that containing excessive “missing” zeros. As a result, it becomes more challenging to distinguish the “true” zeros from the “missing” zeros.

While generating the simulation datasets, the scale factor *β* controls the coverage of simulated contact matrices. We estimated 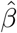 from the published Hi-C data by Rao et al. [6] (Supplementary Methods). At the default 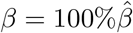 setting, the simulated Hi-C map contained about 4.9 million contacts from the homologous X chromosomes. To evaluate the performance of our methods on lower-coverage data, we fixed the SNP density 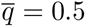 and gradually decreased the value of *β* from 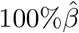 to 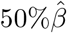, and 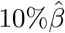, resulting in 2.5 million, 1.0 million, and 0.5 million contacts, respectively. We then repeated the assessments of the ASHIC-ZIPM, ASHIC-PM, mate-rescue, and allele-certain methods with these low-coverage simulation datasets.

As shown in Figure 3A, ASHIC-ZIPM and ASHIC-PM maintained the highest SCC values across all coverage levels. When the sequencing coverage decreased from 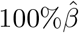 to 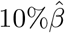, the SCC values for both methods only dropped by 0.28%. On the other hand, when sequencing coverage lowered, the SCC values decreased evidently for mate-rescue and allele-certain by 1.80% and 10.38%, respectively. These results suggested that our ASHIC methods can robustly and accurately infer allele-specific contact matrices under low sequencing coverage conditions.

**Figure 3.**
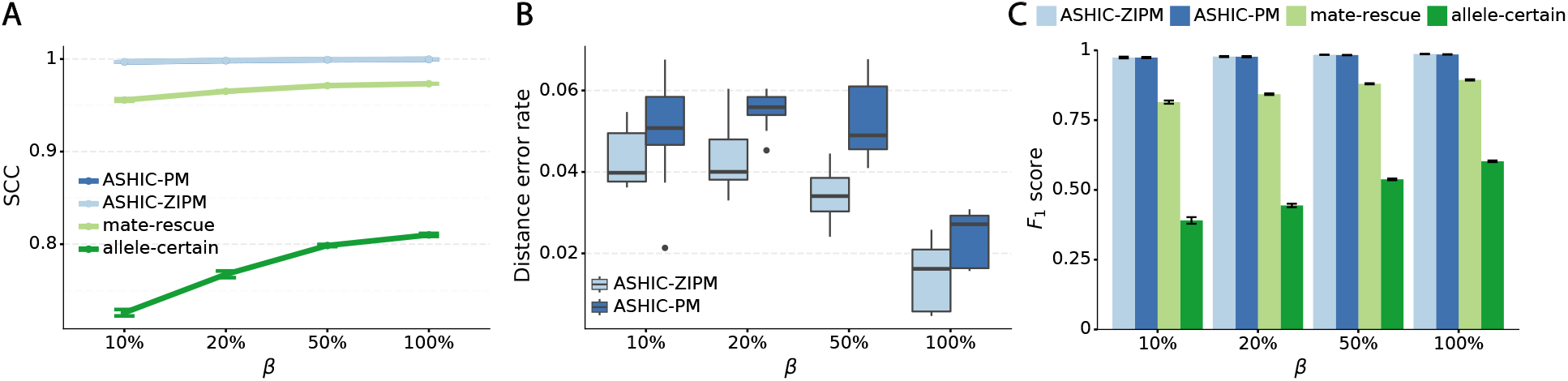
ASHIC-ZIPM accurately imputes diploid contact maps and 3D structures on low-coverage Xa/Xi simulation data. **(A)** SCCs between the imputed diploid contact matrices and the true contact matrices, **(B)** Distance error rates between the predicted allelic 3D structures and the true structures, and **(C)** *F_1_* scores of the identified allele-specific chromatin interactions at different sequencing coverage β levels.

Additionally, we observed that ASHIC-ZIPM produced more accurate 3D structures with smaller distance error rates than those produced by ASHIC-PM across all sequencing coverage levels (Figure 3B). The improvements of the distance error rates were significant at coverage levels 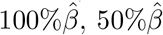 and 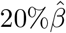 (p-values = 2.53 × 10^−3^,2.53 × 10^−3^,6.26 × 10^−3^, respectively, one-sided paired Wilcoxon signed-rank tests).

When the sequencing coverage decreased from 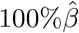 to 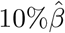, the true set of allele-specific interactions decreased from 9061.5 to 2136.4 interactions (Supplementary Table S5, Supplementary Methods). As shown in Figure 3C, when the coverage decreased from 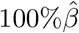 to 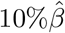, the ability of the allele-certain method to detect allele-specific interactions was highly impacted as its *F*_1_ scores dropped by 35.17% from 0.6024 to 0.3906. The decrease of *F*_1_ score for mate-rescue was less severe, about 8.90% from 0.8940 to 0.8144. The ASHIC methods consistently delivered robust results against coverage changes (ASHIC-ZIPM: Δ*F*_1_ = 1.26%, ASHIC-PM: Δ*F*_1_ = 1.14%), and maintained high *F*_1_ score even at the lowest 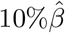 level (ASHIC-ZIPM: 0.9743, ASHIC-PM: 0.9740). The decay in *F*_1_ scores for the allele-certain and mate-rescue methods was primarily contributed by their low recall rates (Supplementary Figure S1).

Taken together, our results demonstrated that the ASHIC methods significantly outperformed other methods in low sequencing coverage conditions, resulted in more accurately imputed matrices and benefited the detection of allele-specific interactions on low-coverage data. In particular, we observed that ASHIC-ZIPM had better performance than ASHIC-PM under low coverage conditions. This is owing to the fact that in our ASHIC-ZIPM model, the Poisson state probabilities ***γ*** act as weights between the “true” and “missing” zeros. When the sequencing coverage lowered, the observed diploid matrices contained additional “missing” zeros. The zero-inflated model explicitly adjusted the estimation of ***γ*** to model these “missing” zeros, thereby achieving better model fitting results. Consistent with our expectations, the estimated values of ***γ*** became smaller as coverage decreased, which demonstrated its ability to account for the additional “missing” zeros (Supplementary Figure S2).

#### 3.1.3 Performance on low SNP density data

In addition to the sequencing coverage, the SNP density is another major factor affecting the performance of the diploid Hi-C methods. The SNP density varies across different species and cell lines. For example, the F1 mouse cross (BL6×Spretus) has a relatively high SNP density of approximately 1 SNP per 75 bp. On average, a 70-bp sequence read has a 60% chance overlapping with SNP(s), thus being allele-identifiable. Whereas the GM12878 cell line has a low SNP density about 1 for every 1700 bp, which is corresponding to an average allele-identifiable probability of 0.04 (Supplementary Table S1). To evaluate the performance of our methods on low-SNP-density data, we fixed the coverage level at 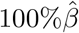 and then gradually decreased 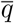, the average allele-identifiable probability, from 0.5 which mimics the BL6×Spretus cross, to 0.25, 0.1, and 0.05, where the smallest value mimics the GM12878 cells.

When the SNP density was low, fewer both-end allele-certain contacts but higher number of one-end allele-ambiguous and both-end allele-ambiguous contacts were observed. Consequently, as the average allele-identifiable probability 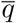 decreased from 0.5 to 0.05, the recovery rates dropped dramatically from 25.65% to 0.25% for allele-certain and from 75.55% to 9.82% for mate-rescue (Supplementary Table S3). In contrast, our ASHIC methods were able to recover all allele-ambiguous reads at the lowest 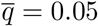 setting. Among the recovered contacts, 15.95% for ASHIC-ZIPM and 17.60% for ASHIC-PM were incorrectly imputed (Supplementary Table S4).

Consistent with the high recovery rates and low imputation error rates, the SCC values also demonstrated robust and accurate imputation of diploid contact matrices by the ASHIC methods at low SNP density settings (Figure 4A). When the average allele-identifiable probability 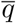 decreased from 0.5 to 0.05, the SCC values dropped significantly from 0.8100 to 0.3959 for allele-certain and from 0.9733 to 0.8719 for mate-rescue, respectively. In contrast, the SCC values remained high at 0.9941 and 0.9922 for ASHIC-ZIPM and ASHIC-PM, respectively, at the lowest 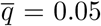 setting. Moreover, ASHIC-ZIPM significantly outperformed ASHIC-PM at the lowest SNP density level (*p*-value = 8.30 × 10^−3^, one-sided paired Wilcoxon signed-rank test). The difference between our ASHIC methods and other methods was also observed on the PCC plot at the lowest SNP density, particularly for long genomic distances (Supplementary Figure S3). Furthermore, when comparing the predicted allelic 3D structures with the ground truth, ASHIC-ZIPM outperformed ASHIC-PM significantly at all SNP density levels (*p*-values = 2.53 × 10^−3^,4.67 × 10^−3^,3.46 × 10^−3^,2.53 × 10^−3^, for 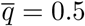, 0.25, 0.1, 0.05, respectively, one-sided paired Wilcoxon signed-rank tests) (Figure 4B).

**Figure 4:**
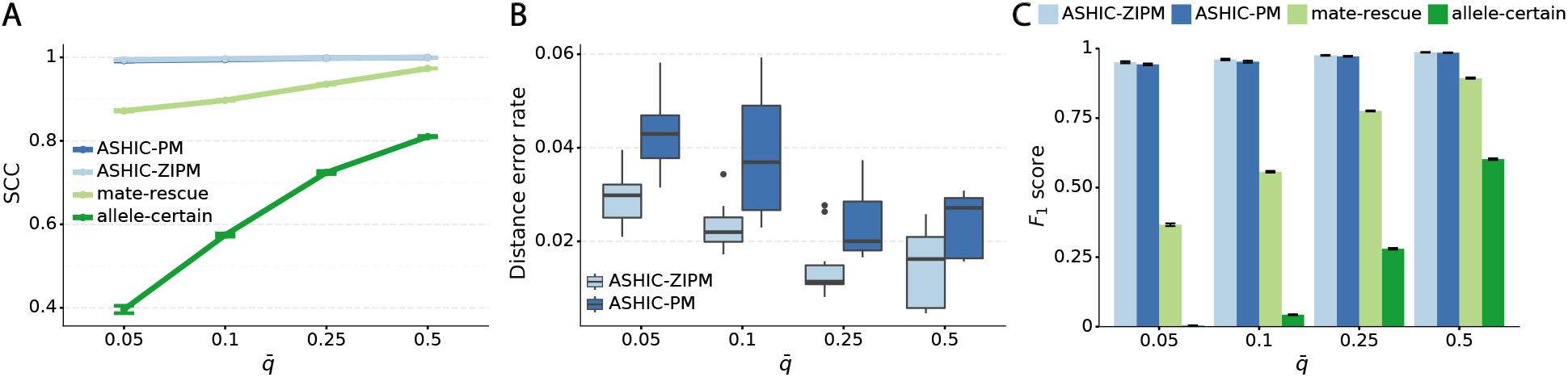
ASHIC-ZIPM accurately imputes diploid contact maps and 3D structures on low-SNP-density Xa/Xi simulation data. **(A)** SCCs between the imputed diploid contact matrices and the true contact matrices, **(B)** Distance error rates between the predicted allelic 3D structures and the true structures, and **(C)** *F*_1_ scores of the identified allele-specific chromatin interactions at different SNP density *q* levels.

Next, we questioned whether the ability to detect allele-specific chromatin interactions was impacted by low SNP density levels. Adjusting the average allele-identifiable probability did not affect the underlying true diploid contact matrices. As a result, the true set of allele-specific interactions remained the same at different SNP density settings (Supplementary Table S5, 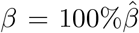). As shown in Figure 4C, low SNP density severely impacted the allele-certain and mate-rescue methods. The *F*_1_ scores of allele-certain dropped from 0.6024 to 0.0039, recovering only 17.8 out of 9061.5 true allele-specific interactions. Similarly, the *F*_1_ score of mate-rescue dropped from 0.8940 to 0.3666. In contrast, when SNP density lowered, the *F*_1_ score of our methods decreased only slightly—3.62% for ASHIC-ZIPM and 4.26% for ASHIC-PM. In addition, our ASHIC methods outperformed the other methods by a notable margin. We observed that decreasing SNP density increased the margin between ASHIC-ZIPM and other methods. Taken together, our results demonstrated that the ASHIC-ZIPM method significantly exceeded other methods with high robustness in low SNP density situations.

### 3.2 Simulation studies on identical chromosomal structures

In the aforementioned simulation settings, we took the homologous X chromosomes in GM12878 cells as the ground truth, where Xa and Xi have drastically dissimilar structures. Unlike the X chromosomes, homologous autosomes often have similar 3D shapes. Imputing diploid Hi-C contact matrices and allelic structures from homologs with similar structures is a more challenging problem than the one from homologs with different structures. To evaluate our methods in such situation, we duplicated the paternal/Xi structure as the pseudo-maternal structure to build an identical homologous structure pair (see Methods). We then generated simulation datasets and evaluated our methods at different coverage and SNP density settings, similarly as previously described.

#### 3.2.1 Performance on low sequencing coverage data

As demonstrated in previous homologous structure simulations, our ASHIC methods maintained high accuracy of imputed diploid contact matrices at low sequencing coverage settings (Figure 5A). The SCC values were all above 0.9949 for ASHIC-ZIPM and above 0.9938 for ASHIC-PM at various sequencing coverage levels. On the other hand, the SCC values of mate-rescue demonstrated a minor decline from 0.9778 to 0.9664 when the coverage decreased from 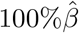 to 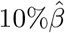. The allele-certain method was the most impacted, as its SCC values declined by 7.46% from 0.8362 to 0.7738 when the coverage level dropped from 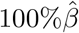 to 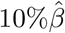.

We then evaluated the accuracy of the allelic 3D structures predicted by our ASHIC methods. Overall, ASHIC-ZIPM generated more accurate structures with smaller distance error rates than the ones predicted by ASHIC-PM across all coverage levels (Figure 5B). The improvements were significant at 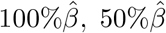, and 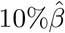 levels (*p*-values = 2.53 × 10^−3^, 1.42 × 10^−2^, 2.53 × 10^−3^, one-sided paired Wilcoxon signed-rank tests).

In addition to comparing the predicted allelic structures against the ground truth structures, we further calculated the homologous distance error rate between the predicted maternal and paternal structures (Figure 5C). For both ASHIC-ZIPM and ASHIC-PM methods, the average homologous distance error rates were smaller than 0.08, suggesting that both models produced homologous structures with very similar shapes. Furthermore, the ASHIC-ZIPM model had significantly lower homologous distance error rates than ASHIC-PM, at sequencing coverage 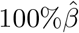 and 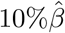 levels (*p*-values = 2.53 × 10^−3^, one-sided paired Wilcoxon signed-rank tests). These results further confirmed that ASHIC-ZIPM predicted more accurate allelic 3D structures than the structures predicted by ASHIC-PM.

Next, we investigated the effects of low sequencing coverage on the detection of chromatin interactions when the homologous structures were identical. Similar to the case of different homologous structures, we applied Fit-Hi-C [23] to call significant interactions on the two allele-specific contact matrices separately. Given that the two ground truth homologous structures were identical, we defined the true integration set as the bi-allelic interactions shared by both maternal and paternal chromosomes (Supplementary Methods). When the coverage dropped from 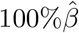 to 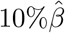, the number of interactions in the true set decreased by 81.48% from 4103.1 to 759.9 (Supplementary Table S6). As shown in Figure 5D, the allele-certain method was the most impacted by the sequencing coverage changes, where its *F*_1_ scores decreased by 31.23% from 0.6127 to 0.4214 as the coverage dropped from 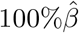 to 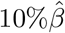. The *F*_1_ score of mate-rescue decreased to a less extend, by 7.37% from 0.9075 to 0.8406. Whereas our ASHIC-ZIPM and ASHIC-PM methods demonstrated consistent high *F*_1_ scores of 0.9351 and 0.9296, respectively, even under the lowest coverage 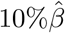 setting.

#### 3.2.2 Performance on low SNP density data

When the SNP density lowered, we observed an overall decreasing trend in the SCC values for all four methods (Figure 6A). The allele-certain and mate-rescue methods were greatly impacted by the low SNP density. When the average allele-identifiable probability 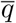 decreased from 0.5 to 0.05, the SCC values dropped significantly by 46.52% from 0.8362 to 0.4472 for allele-certain and by 9.16% from 0.9778 to 0.8883 for mate-rescue. Again, our ASHIC methods maintained robustly high accuracy of the imputed contact matrices; the SCC values decreased only by 0.45% from 0.9996 to 0.9950 for ASHIC-ZIPM and by 2.38% from 0.9988 to 0.9750 for ASHIC-PM when 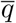 decreased from 0.5 to 0.05. The visible difference between ASHIC-ZIPM and ASHIC-PM at the lowest SNP density level 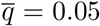 was also supported by the PCC measures, where ASHIC-ZIPM outperformed ASHIC-PM by an evidently large margin of PCCs within genomic distance of 100 Mb (Supplementary Figure S3).

In terms of structural accuracy, ASHIC-ZIPM also outperformed ASHIC-PM with significantly smaller distance error rates across all SNP density levels (*p*-values = 2.53 × 10^−3^, one-sided paired Wilcoxon signed-rank tests) (Figure 6B). Furthermore, the allelic structures predicted by ASHIC-ZIPM demonstrated significantly smaller homologous distance error rates than the ones predicted by ASHIC-PM (*p*-values = 2.53 × 10^−3^, at all four 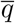 levels, one-sided paired Wilcoxon signed-rank tests) (Figure 6C).

In addition to achieving the highest imputation accuracy of the diploid contact matrices and 3D structures, ASHIC-ZIPM also demonstrated the best performance with respect to the detection of biallelic chromatin interactions under low SNP density conditions (Figure 6D). When the average allele-identifiable probability 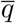 decreased from 0.5 to 0.05, the *F*_1_ values dropped by 99.43% for allele-certain, 62.49% for mate-rescue, and 8.32% for ASHIC-PM. The ASHIC-ZIPM model showed the smallest decline in *F*_1_ scores, merely 1.70% from 0.9814 to 0.9647. Moreover, we observed that ASHIC-ZIPM significantly outperformed all other methods by a large margin across all SNP density levels (*p*-values = 2.53 × 10^−3^, one-sided paired Wilcoxon signed-rank tests)

Taken together, we demonstrated that our ASHIC methods significantly outperformed the allele-certain and mate-rescue methods under low SNP density conditions when the homologous structures have identical shapes. In addition, ASHIC-ZIPM evidently outperformed the ASHIC-PM model by a large margin, especially at the lowest SNP density level.

### 3.3 ASHIC-ZIPM reconstructs the bipartite structure of inactive X chromosome

The X chromosomes in mammalian females is a representative example of homologous structural difference. In contrast to males with only one X chromosome, the females have two copies of X chromosome. To compensate for the dosage imbalance of X-linked genes between females and males, one X chromosomes in female cells is randomly silenced through the X chromosome inactivation (XCI) mechanism [24]. To study the structural differences between the active X (Xa) and inactive X (Xi) chromosomes, we applied our ASHIC-ZIPM model to a published diploid Hi-C data generated from wild-type patski (BL6× *Spretus*) cells [15]. The patski cell line has completely skewed XCI such that the maternal BL6 X is always inactive while the paternal *Spretus* X is always active. Several Hi-C studies conducted on the patski cells have demonstrated that the maternal Xi and paternal Xa chromosomes exhibit distinct morphology and chromatin contact profiles [9, 15]. Specifically, Xi shows a clear bipartite structure, where the entire chromosome is densely packed into two superdomains. The hinge region (chrX:75,637,519-75,764,753 bp) between the two superdomains contains the macrosatellite repeat locus *Dxz4* and represents a nucleolus-associated domain [6, 9, 11, 15].

To study the bipartite organization of Xi, we applied our ASHIC-ZIPM model to the patski Hi-C data and reconstructed the diploid contact maps and 3D structures of Xa and Xi at various resolutions (1 Mb, 500 kb, and 100 kb). As shown in Figure 7A, the contact map of Xa demonstrated a clear plaid pattern representing the alternating A/B compartments. In contrast, Xi was clearly separated into two superdomains by a hinge region containing *Dxz4*. We observed frequent intra-superdomain contacts but sparse inter-superdomain contacts on Xi. In addition, we calculated the bipartite index (BI) [9] (Supplementary Methods) for both X chromosomes (Figure 7B). At all three resolutions, we observed an evident BI peak at the hinge region (*Dxz4*) on Xi, confirming the existence of bipartite organization on Xi. In contrast, the BI values were rather flat across the entire Xa, indicating the absence of bipartite structure. These observations demonstrated that our ASHIC-ZIPM method can produce robust and consistent diploid contact maps across different resolutions.

**Figure 5:** ASHIC-ZIPM accurately imputes diploid contact maps and 3D structures on low-coverage identical-homolog simulation data. **(A)** SCCs between the imputed diploid contact matrices and the true contact matrices, **(B)** Distance error rates between the predicted allelic 3D structures and the true structures, **(C)** Homologous distance error rates between the predicted maternal and paternal 3D structures, and **(D)** *F*_1_ scores of the identified bi-allelic interactions at various sequencing coverage *β* levels.

**Figure 6:** ASHIC-ZIPM accurately imputes diploid contact maps and 3D structures on low-SNP-density identical-homolog simulation data. **(A)** SCCs between the imputed diploid contact matrices and the true contact matrices, **(B)** Distance error rates between the predicted allelic 3D structures and the true structures, **(C)** Homologous distance error rates between the predicted maternal and paternal 3D structures, and **(D)** *F*_1_ scores of the identified bi-allelic interactions at different SNP density *q* levels.

**Figure 7:**
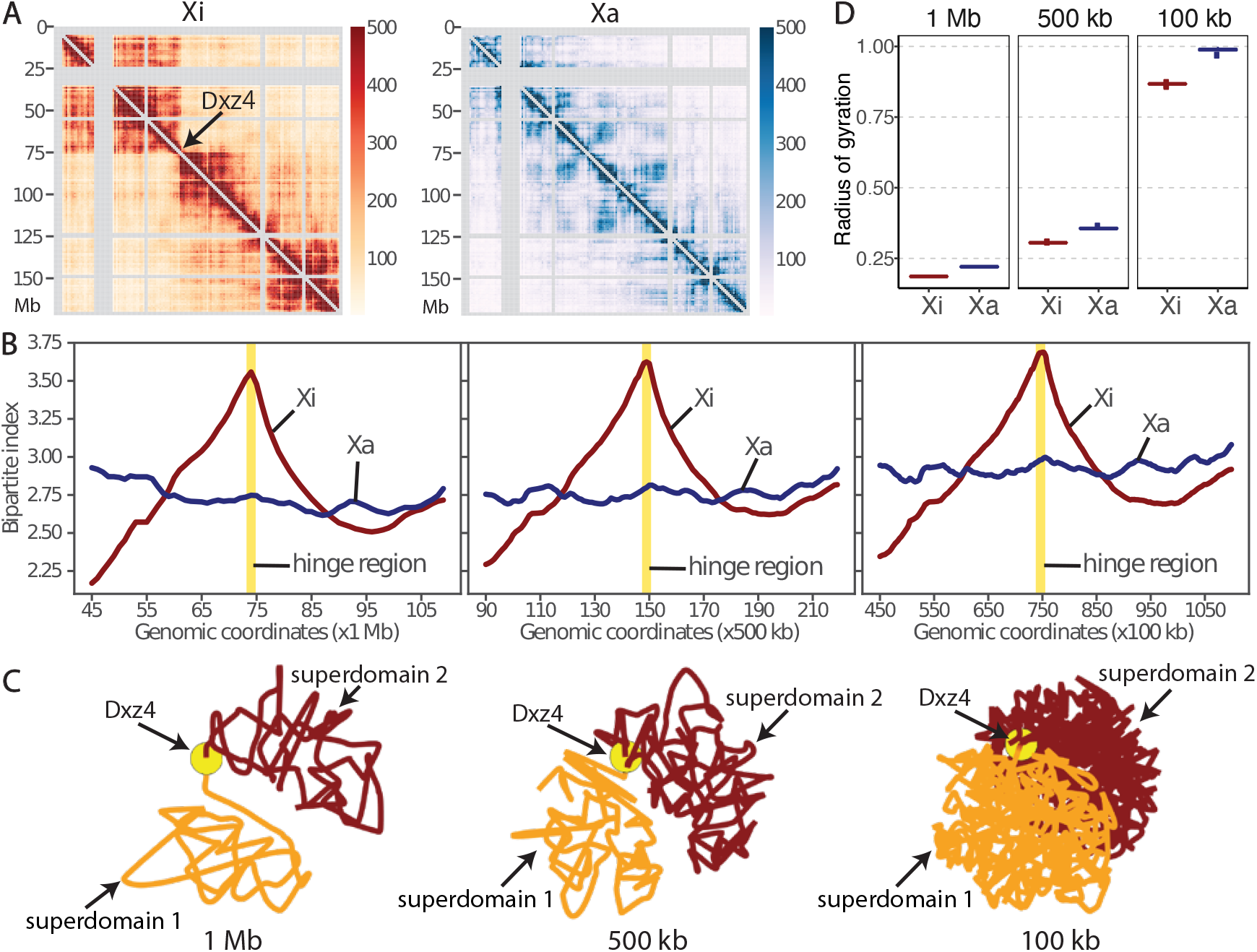
Bipartite organization of the inactive X chromosome in mouse patski cells. **(A)** ASHIC-ZIPM-imputed allele-specific Hi-C contact matrices of Xi and Xa are shown at 1 Mb resolution. The Xi shows a bipartite structure of two superdomains connected by a hinge region (*Dxz4*), indicated by an arrow. Gray strips indicate low mappability regions. **(B)** Chromosomewise bipartite index (BI) values for Xi (brown) and Xa (blue) at 1 Mb (left), 500 kb (middle), and 100 kb (right) resolutions. The Xi curve shows an evident peak at the hinge region (yellow). **(C)** The Xi structures predicted by ASHIC-ZIPM at 1 Mb, 500 kb, and 100 kb resolutions. The first superdomain (centromeric region) is shown in orange, and the second superdomain (distal region) is shown in brown. The hinge region (*Dxz4*) is marked by a yellow ball. The 3D structures are interpolated and smoothed by the Akima interpolator in `SciPy`. **(D)** Box plots of the radius of gyration for the Xi (brown) and Xa (blue) structures at 1 Mb, 500 kb, and 100 kb resolutions.

In addition to the existence of two superdomains in the Xi contact map, we also observed that the predicted Xi structures preserved the bipartite conformation across all three resolutions (Figure 7C). The two superdomains were clearly separated in space, as each superdomain occupied half of the sphere and there were minimal interactions between them. In addition, the hinge region (*Dxz4*) connecting the two superdomains was located towards the periphery of the Xi structure, which is consistent with previous DNA-FISH results [9]. While the previously published Xa and Xi structures were at 1 Mb [9] and 500 kb [14] resolutions, our method produced chromosomal structures at 100 kb resolution and successfully confirmed the bipartite organization of Xi.

With regards to the overall morphology of the chromosomal structures, we observed that Xi exhibited a more condensed structure than Xa, which is consistent with the fact that Xi is almost entirely silenced. In particular, we calculated the radius of gyration (*R_g_*, Supplementary Methods) [25] to measure the compactness of the X chromosomes (Figure 7D). Across all three resolutions, Xi consistently showed a significantly lower *R_g_* value than Xa, indicating that Xi was more tightly packed (*p*-values = 4.43 × 10^−5^, one-sided paired Wilcoxon signed-rank tests).

Collectively, the results obtained on the published patski Hi-C data demonstrated that our ASHIC-ZIPM method can accurately detect distinct allele-specific chromatin organizations of Xa and Xi at fine resolution.

### 3.4 ASHIC-ZIPM reveals differential allele-specific chromatin organization at the *H19/Igf2* imprinting regions

Imprinting is an epigenetic mechanism that causes a subset of genes to express exclusively on one allele in diploid cells. The expression of imprinted genes is controlled by parental-specific epigenetic modifications, such as DNA methylation, at the imprinting control regions. One well-studied example is the *H19/Igf2* imprinting region. In the mouse genome, the paternally expressed *Igf2* gene is located approximately 80 kb upstream (telomeric side) from the long non-coding RNA *H19* that is expressed only on the maternal allele. These two genes demonstrate opposite allelespecific expression yet share a common set of enhancers located downstream of *H19* [27–29]. It has been shown that the parent-specific expression pattern of *H19* and *Igf2* is controlled by the H19 differentially methylated region (H19-DMR) located 2 kb upstream from *H19* [30]. The H19-DMR is methylated only on the paternal allele, and therefore exhibits methylation-sensitive CCCTC-binding factor (CTCF) binding. On the maternal allele, the unmethylated H19-DMR recruits CTCF bindings and therefore blocking the interactions between the enhancers and *Igf2*. As a result, *Igf2* remains unexpressed, while *H19* can still access the enhancers and thus is activated. Whereas on the paternal allele, the methylated H19-DMR inhibits CTCF bindings. Consequently, *Igf2* can access the enhancers and being activated; while the *H19* silencing is likely caused by spreading of methylation from H19-DMR [31].

It has been widely speculated that CTCF attains enhancer-blocking insulation function via the formation of chromatin loops [32]. Using diploid Hi-C contact maps of human GM12878 cells at 25 kb resolution, Rao et al. [6] examined the *H19/IGF2* imprinting region and identified parental-specific chromatin loops between the *H19/IGF2* cluster and a distal region which was referred to as the H19/Igf2 Distal Anchor Domain (HIDAD). The HIDAD-H19 loop was present exclusively on maternal allele; in contrast, the HIDAD-*IGF2* loop appeared only on the paternal allele. Additionally, Lléres et al. [33] performed a diploid 4C-seq study on the mouse ESCs and showed that H19-DMR interacted significantly more with the mouse homologue of HIDAD (mHIDAD) on maternal allele compared to the interactions on the paternal allele. They subsequently performed 3D DNA-FISH experiments and confirmed that the distances between mHIDAD and *H19* were significantly shorter on the maternal allele than the distances on the paternal allele.

Although the aforementioned 4C-seq study [33] and several other 3C studies [34–36] have been conducted in the *H19/Igf2* imprinting region, diploid Hi-C studies are still restricted to a rather coarse resolution due to the limitations of low SNP density and insufficient sequencing coverage. To bridge this gap and provide a holistic view of chromatin structures on the imprinted *H19/Igf2* region, we applied our ASHIC-ZIPM method to the published diploid Hi-C data in mouse patski cells [15], and generated fine-scale allele-specific contact maps and 3D structures of a 5-Mb region (chr7: 140—145 Mbp) around the *H19/Igf2* imprinting region at 10 kb resolution.

First, we constructed a differential contact map using log-fold-change values between the imputed maternal and paternal contacts (Figure 8A). Along with the contact map, we also visualized the allelic CTCF ChIP-seq data [15]. Consistent with previous studies [37, 38], we observed a clear maternal-specific CTCF binding at the H19-DMR locus. Additionally, a few bi-allelic CTCF binding clusters were observed at mHIDAD, near the *Syt8* and *Lsp1* genes, and at the telomeric side of *Igf2.* As shown in Figure 8A, the contacts between mHIDAD and *H19* were enriched on the maternal allele (box 1), whereas the contacts between mHIDAD and *Igf2* were enriched on the paternal allele (box 2). In addition to the contacts between mHIDAD and *H19/Igf2, H19* and *Igf2* demonstrated differential contact preferences to the bi-allelic CTCF clusters near *Syt8* and *Lsp1* (boxes 3 and 4). To further characterize the parental-specific chromatin interactions, we identified chromatin loops with genomic distance of 30–500 kb from the two imputed allelic contact maps using Fit-Hi-C [23] with a strict FDR threshold (*q*-value < 10^−5^). The identified chromatin loops were mostly anchored to the CTCF binding clusters (Figure 8A). We further categorized these chromatin loops into bi-allelic loops that were shared between the two alleles, or monoallelic loops that are either maternal-specific or paternal-specific. Consistent with the differential contact map, chromatin loops anchored at *H19* and *Igf2* were primarily parental-specific. We observed a distinct pattern of maternal-specific chromatin loops between mHIDAD and *H19* and paternal-specific chromatin loops between mHIDAD and *Igf2*. Besides mHIDAD, the region containing bi-allelic CTCF binding clusters near the *Syt8* and *Lsp1* genes also demonstrated parental-specific chromatin interactions with *H19* and *Igf2*. Specifically, these CTCF clusters interacted preferentially with *H19* on the maternal allele and with *Igf2* on the paternal allele. These observations are consistent with the previous 4C-seq results in mouse ESCs [33].

**Figure 8:**
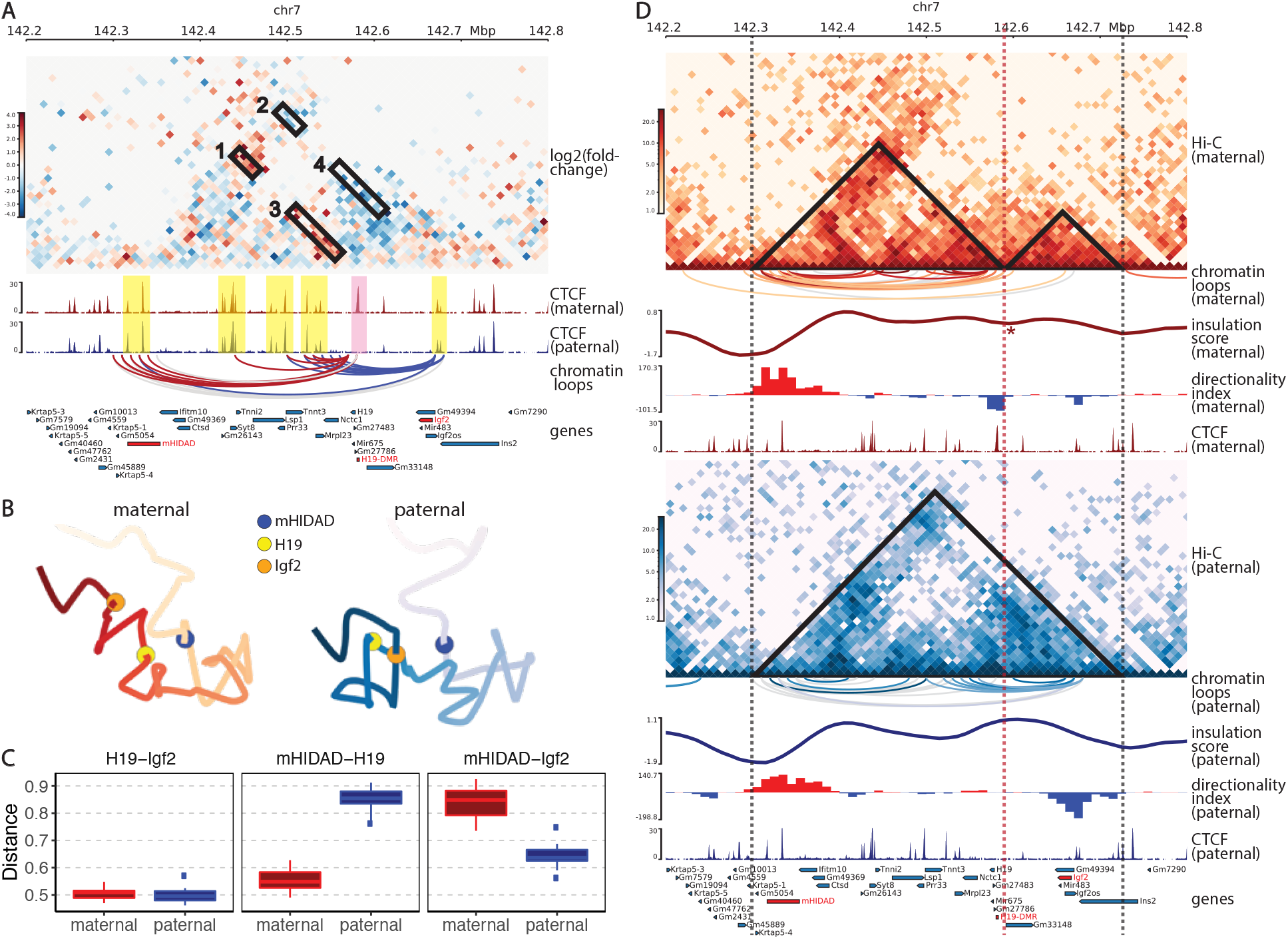
Allele-specific chromatin organizations of the *H19/Igf2* imprinting region in mouse patski cells. **(A)** Differential contact map between the ASHIC-ZIPM-imputed maternal and paternal contacts at 10 kb resolution. Contact counts are normalized separately on each allele to account for the potential mapping bias towards the reference genome. The red vs blue color key indicates maternal vs paternal enrichment. Four allelicly enriched chromatin interacting regions are labeled in boxes 1–4. Maternal-specific CTCF peak (pink) and bi-allelic CTCF binding clusters (yellow) are highlighted. Chromatin loops are called using Fit-Hi-C [23] and categorized into maternal-specific (red), paternal-specific (blue), and bi-allelic (gray). Only loops anchored at *H19* or *Igf2* are displayed. **(B)** Allelic 3D structures of the *H19/Igf2* imprinting region predicted by ASHIC-ZIPM at 10 kb resolution. The maternal (red) and paternal (blue) structures are overall similar, but the relative spatial positions of mHIDAD (blue), *H19* (yellow), and *Igf2* (orange) are evidently different. **(C)** Box plots of pairwise Euclidean distances between *H19-Igf2* (left), mHIDAD-*H19* (middle), and mHIDAD-*Igf2* (right). **(D)** Allelic Hi-C contact maps at 10 kb resolution (top panel: maternal allele, red color key; bottom panel: paternal allele, blue color key). Maternal-specific (red), paternal-specific (blue), and bi-allelic (gray) chromatin loops are called using Fit-Hi-C [23]. A local minimum of the insulation score (IS) is marked by an asterisk. Positive and negative directionality index (DI) values [8] are shown in red and blue, respectively. (Sub-)TAD domains derived from IS and DI measures are labeled as triangles on the contact maps, and dashed lines indicate (sub-)TAD boundaries. Panels (A) and (D) are drawn using `pyGenomeTracks` [26].

Besides the differential contact map, we also examined the allele-specific chromatin conformations using the predicted allelic 3D structures (Figure 8B). The overall chromatin organizations of the *H19/Igf2* imprinting region appeared to be similar between the two alleles. However, the relative spatial position among mHIDAD, *H19*, and *Igf2* demonstrated parental-specific differences. From the 3D structures, we observed that mHIDAD was spatially close to *H19* on the maternal allele, presumably forming a chromatin loop. In addition, we observed that *Igf2* was much closer to mHIDAD on paternal structure than on the maternal structure.

For the quantitative comparison, we calculated the pairwise Euclidean distances of mHIDAD, *H19*, and *Igf2* on the maternal and paternal structures predicted by ASHIC-ZIPM from 20 random initializations. As shown in Figure 8C, the distance between mHIDAD and *H19* was significantly smaller on the maternal structure than that on the paternal structure (*p*-value = 4.43 × 10^−5^, one-sided Wilcoxon paired signed-rank test), which is consistent with the previous DNA-FISH data [33]. In contrast, the distance between mHIDAD and *Igf2* was significantly larger on maternal allele (*p*-value = 4.43 × 10^−5^, one-sided Wilcoxon paired signed-rank test), which is consistent with the observation of paternal-specific HIDAD-IGF2 loop in human GM12878 cells [6]. No significant difference of the distance between *H19* and *Igf2* was detected on our predicted allelic structures. These observations demonstrated that our method can stably predict fine-scale 3D structures that reflect the distinct parental-specific chromatin conformations.

### 3.5 ASHIC-ZIPM-imputed diploid contact maps uncover the maternal-specific sub-TAD organization at imprinted *H19/Igf2* locus

In addition to the formation of chromatin loops, CTCF also participates in the establishment of higher-order chromatin structures such as topologically-associating domains (TADs). TADs are sub-megabase genomic regions containing frequent local chromatin interactions, whereas TAD boundaries result in physical insulation between neighboring domains [8]. It has been observed that CTCF bindings are often enriched at TAD boundaries and play an important role in TAD formation [6, 8]. Since the genome is organized in a hierarchical manner, smaller domains called sub-TADs are often observed within the large TADs. Unlike TADs that are mostly invariant between cell types, sub-TADs are more variable and play a pivotal role in mediating cell-type-specific gene regulation [39, 40]. Based on the presence of monoallelic CTCF bindings at H19-DMR, Llères et al. [33] proposed a novel parental-specific sub-TAD model for the regulation of imprinting at *H19/Igf2* locus. Supported by allelic 4C-seq and DNA-FISH data, they speculated that several bi-allelic CTCF binding sites form a first layer of TAD on both alleles. In addition, the maternal-specific CTCF binding around H19-DMR hijacks the first layer of TAD and consequently creates an additional layer of sub-TAD on the maternal allele.

To verify this hypothesis, we calculated the insulation score (IS) [41] and directionality index (DI) [8] using `TADtool` [42] to search for possible (sub-)TAD boundaries around the *H19/Igf2* imprinting region. Overall we observed similar IS values on both alleles, except at the H19-DMR locus (Figure 8D, Supplementary Figure S4). Specifically, we observed a local minimum of IS values at H19-DMR only on the maternal allele indicating a potential presence of a sub-TAD boundary at H19-DMR. Consistently, the DI values suggested similar (sub-)TAD pattern (Figure 8D). We observed strong positive DIs at mHIDAD on both alleles, indicating that mHIDAD is highly biased towards interacting with its downstream loci and serves as a starting position of a TAD. On the other hand, the telomeric-side flanking region of *Igf2* demonstrated negative DIs on both alleles, indicating a likely ending boundary of a TAD. Furthermore, a negative DI region around H19-DMR appeared only on the maternal allele, suggesting H19-DMR has a higher tendency to interact with its upstream loci, possibly indicating an ending position for a maternal-specific sub-TAD.

Both the IS and DI measurements suggested that *H19/Igf2* is embedded within a TAD demarcated by two main boundaries: one near mHIDAD and the other one at the telomeric side of *Igf2*. The locations of the two boundaries were in good agreement between both alleles. However, the (sub-)TAD organization within this TAD region undergoes drastic parental-specific changes. Specifically, we observed a sub-TAD boundary at H19-DMR locus exclusively on the maternal allele. The TAD and sub-TAD boundaries mentioned above were all located at CTCF binding clusters. We further examined the allelic chromatin loops within this imprinting region (Figure 8D). On the maternal allele, chromatin loops were mostly confined to the mHIDAD-*H19* sub-TAD. Whereas on the paternal allele, we observed several chromatin loops connecting the centromeric side of H19-DMR with *Igf2*, indicating the absence of insulation at H19-DMR. These observations of allelic chromatin loops are consistent with the parental-specific (sub-)TAD structures.

Taken together, these results supported the hypothesis that the maternal-specific CTCF binding at H19-DMR forms a chromatin loop with the CTCF binding sites at mHIDAD. This mHIDAD-*H19* loop creates an additional layer of sub-TAD inside the original mHIDAD-*Igf2* TAD. The maternal-specific mHIDAD-*H19* sub-TAD organization mediates the insulation between the centromeric side of H19-DMR and *Igf2*, and thereby leading to the silencing of *Igf2* on the maternal allele.

## 4 DISCUSSION

In this work, we proposed a hierarchical Bayesian framework for imputing allele-specific contacts in diploid Hi-C data and reconstructing allelic 3D structures. We developed two models under this Bayesian framework: ASHIC-PM and ASHIC-ZIPM. To the best of our knowledge, our ASHIC methods are the first methods that produce fully decomposed diploid Hi-C contact matrices as well as the allelic 3D structures.

Unlike the existing allele-certain and mate-rescue approaches, our ASHIC methods utilize all diploid Hi-C contacts, including both-end allele-ambiguous contacts. As a result, ASHIC methods exceeded the allele-certain and mate-rescue methods, in terms of producing more accurate diploid matrices and structures as well as facilitating better detection of allele-specific chromatin interactions. We also conducted a series of simulation experiments to evaluate how the performance our methods are impacted by various factors, including sequencing coverage, SNP density, and homologous structural similarity. Overall, our models significantly outperformed other methods, especially under low sequencing coverage and low SNP density conditions. The ability of the ASHIC methods in inferring allele-ambiguous contacts at low-SNP-density setting is critical for analyses in diploid human cells such as GM12878, where the existing mate-rescue method [6] was only able to rescue 0.14% of total diploid contacts (Supplementary Table S1).

In our simulation studies, we did not compare the ASHIC methods with the recently published Dip-C method by Tan et al. [13] as their method was specifically designed for single-cell Hi-C data. Another reason was that Dip-C does not impute intra-chromosomal both-end allele-ambiguous contacts. Therefore we expect that its performance would be close to the mate-rescue method. In addition, our earlier work of the Poisson-Gamma model [9] imputes diploid contact counts based on genomic distances rather than the spatial distances derived from 3D structures, and therefore is not computationally stable on fine-resolution (such as 100 kb) or low-coverage Hi-C data. Lastly, the newly developed diploid-PASTIS method by Cauer et al. [14] predicts only the allelic 3D structures rather than the diploid contact matrices. Therefore, we did not evaluate the diploid-PASTIS method in our simulations as most of our evaluation metrics were based on imputed contact matrices.

The main advantage of the ASHIC-ZIPM model over the ASHIC-PM model is that ASHIC-ZIPM explicitly accounts for the excessive zeros in Hi-C matrices, by modeling the probabilities whether each observed zero count is a “true” zero or a “missing” zero. As a result, we observed that the ASHIC-ZIPM model consistently outperformed the ASHIC-PM model in all simulation settings. While the performance of the two models were often similar, the improvements of ASHIC-ZIPM over ASHIC-PM became more evident when the SNP density decreased. In addition, the differences between the ASHIC-ZIPM and ASHIC-PM models were particularly noticeable under the more challenging simulation setting of identical homologous structures. This is owing to the fact that when SNP density was low, only few allele-certain contacts were observed. The ASHIC-PM model uses the allele-certain contacts to initialize the EM algorithm and treats all zeros as “true” zeros, thereby producing less optimal results. In contrast, ASHIC-ZIPM explicitly adjusts the weights between “true” and “missing” zeros and thereby archiving more accurate models.

We demonstrated two applications of our ASHIC-ZIPM method: the Xa/Xi chromosomes and the *H19/Igf2* imprinting region in the mouse patski cells. Previous studies predicted allelic X chromosome structures at 1 Mb [9] and 500 kb [14] resolutions. In contrast, our method utilizes all diploid contacts and produces finer-scale allelic structures of the entire X chromosomes at 100-kb resolution. Our results further confirmed the existence of the bipartite structure of Xi. The ability to impute all allele-ambiguous contacts is particularly important when zooming into local imprinting regions. Since imprinting regions are often small, fine-resolution allelic contact maps and 3D structures are required for an in-depth study. With our ASHIC-ZIPM model, we produced the first 10-kb-resolution diploid Hi-C contact maps of the *H19/Igf2* imprinting region, and revealed the existence of the maternal-specific sub-TAD organization at H19-DMR. This sub-TAD formation creates an insulation between *H19* and *Igf2* that likely prevents the activation of *Igf2* on the maternal allele. Furthermore, the diploid Hi-C maps produced by ASHIC-ZIPM offer an informative view of the (sub-)TAD organizations on the imprinting region, whereas the previous 4C-seq study [33] were restricted to only few anchor regions.

Currently, only a few limitations can be attributed to our ASHIC methods. First, our methods provide chromosome-wide modeling of diploid Hi-C data. One possible future extension is to build a genome-wide model by incorporating an additional estimation step in the EM algorithm to model the relative position of multiple homologous chromosomes. We could further parallelize the optimization procedures for each homologous chromosome pair to speed up the genome-wide modeling. Second, our model is specifically designed for diploid genomes. Extending our model to polyploid or aneuploid genomes is beyond the scope of this paper. Lastly, the computational efficiency of our EM algorithm, especially the structure estimation step, could be further improved. One possible solution is to adapt an iterative modeling strategy similar to [13, 25], starting with coarse-resolution modeling and gradually refining the structures to finer resolutions.

## 5 ACKNOWLEDGEMENTS

This work is supported by Grant DBI-1751317 from the National Science Foundation, and the Regents’ Faculty Fellowship from the University of California Riverside.

## Conflict of interest

None declared.

## Notes

### Competing Interest Statement

The authors have declared no competing interest.

## Reference

[1] J. Dekker. Gene regulation in the third dimension. Science, 319(5871):1793–1794, 2008.

[2] E. Lieberman-Aiden, N. L. van Berkum, L. Williams, M. Imakaev, T. Ragoczy, A. Telling, I. Amit, B. R. Lajoie, P. J. Sabo, M. O. Dorschner, R. Sandstrom, B. Bernstein, M. A. Bender, M. Groudine, A. Gnirke, J. Stamatoyannopoulos, L. A. Mirny, E. S. Lander, and J. Dekker. Comprehensive mapping of long-range interactions reveals folding principles of the human genome. Science, 326(5950):289–293, 2009.

[3] Z. Duan, M. Andronescu, K. Schutz, S. McIlwain, Y. J. Kim, C. Lee, J. Shendure, S. Fields, C. A. Blau, and W. S. Noble. A three-dimensional model of the yeast genome. Nature, 465(7296):363–367, 2010.

[4] R. Kalhor, H. Tjong, N. Jayathilaka, F. Alber, and L. Chen. Genome architectures revealed by tethered chromosome conformation capture and population-based modeling. Nature biotechnology, 30(1):90–98, 2012.

[5] W. Ma, F. Ay, C. Lee, G. Gulsoy, X. Deng, S. Cook, J. Hesson, C. Cavanaugh, C. B. Ware, A. Krumm, J. Shendure, C. A. Blau, C. M. Disteche, W. S. Noble, and Z. Duan. Fine-scale chromatin interaction maps reveal the cis-regulatory landscape of lincRNA genes in human cells. Nature methods, 12(1):71–78, 2015.

[6] S. S. P. Rao, M. H. Huntley, N. Durand, C. Neva, E. K. Stamenova, I. D. Bochkov, J. T. Robinson, A. L. Sanborn, I. Machol, A. D. Omer, E. S. Lander, and E. L. Aiden. A 3D map of the human genome at kilobase resolution reveals principles of chromatin looping. Cell, 159(7):1665–1680, 2014.

[7] V. Ramani, D. A. Cusanovich, R. J. Hause, W. Ma, R. Qiu, X. Deng, C. A. Blau, C. M. Disteche, W. S. Noble, J. Shendure, and Z. Duan. Mapping 3D genome architecture through in situ DNase Hi-C. Nature protocols, 11(11):2104–2121, 2016.

[8] J. R. Dixon, S. Selvaraj, F. Yue, A. Kim, Y. Li, Y. Shen, M. Hu, J. S. Liu, and B. Ren. Topological domains in mammalian genomes identified by analysis of chromatin interactions. Nature, 485(7398):376–380, 2012.

[9] X. Deng, W. Ma, V. Ramani, A. Hill, F. Yang, F. Ay, J. B. Berletch, C. A. Blau, J. Shendure, Z. Duan, W. S. Noble, and C. M. Disteche. Bipartite structure of the inactive mouse X chromosome. Genome biology, 16(1):152, 2015.

[10] L. Giorgetti, B. R. Lajoie, A. C. Carter, M. Attia, Y. Zhan, J. Xu, C. J. Chen, N. Kaplan, H. Y. Chang, E. Heard, and J. Dekker. Structural organization of the inactive X chromosome in the mouse. Nature, 535(7613):575–579, 2016.

[11] E. M. Darrow, M. H. Huntley, O. Dudchenko, E. K. Stamenova, N. C. Durand, Z. Sun, S. C. Huang, A. L. Sanborn, I. Machol, M. Shamim, A. P. Seberg, E. S. Lander, B. P. Chadwick, and E. L. Aiden. Deletion of DXZ4 on the human inactive X chromosome alters higher-order genome architecture. Proceedings of the National Academy of Sciences, 113(31):E4504–E4512, 2016.

[12] Z. Du, H. Zheng, B. Huang, R. Ma, J. Wu, X. Zhang, J. He, Y. Xiang, Q. Wang, Y. Li, J. Ma, X. Zhang, K. Zhang, Y. Wang, M. Q. Zhang, J. Gao, J. R. Dixon, X. Wang, J. Zeng, and W. Xie. Allelic reprogramming of 3D chromatin architecture during early mammalian development. Nature, 547(7662):232–235, 2017.

[13] L. Tan, D. Xing, C. H. Chang, H. Li, and X. S. Xie. Three-dimensional genome structures of single diploid human cells. Science, 361(6405):924–928, 2018.

[14] A. G. Cauer, G. Yardımcı, J. P. Vert, N. Varoquaux, and W. S. Noble. Inferring diploid 3D chromatin structures from Hi-C data. In 19th International Workshop on Algorithms in Bioinformatics (WABI 2019). Schloss Dagstuhl-Leibniz-Zentrum fuer Informatik, 2019.

[15] G. Bonora, X. Deng, H. Fang, V. Ramani, R. Qiu, J. B. Berletch, G. N. Filippova, Z. Duan, J. Shendure, W. S. Noble, and C. M. Disteche. Orientation-dependent Dxz4 contacts shape the 3D structure of the inactive X chromosome. Nature communications, 9(1):1445, 2018.

[16] A. P. Dempster, N. M. Laird, and D. B. Rubin. Maximum likelihood from incomplete data via the EM algorithm. Journal of the Royal Statistical Society: Series B (Methodological), 39(1):1–22, 1977.

[17] N. Varoquaux, F. Ay, W. S. Noble, and J. P. Vert. A statistical approach for inferring the 3D structure of the genome. Bioinformatics, 30(12):i26–i33, 2014.

[18] S. Wang, J. Xu, and J. Zeng. Inferential modeling of 3D chromatin structure. Nucleic acids research, 43(8):e54–e54, 2015.

[19] J. Dekker, M. A. Marti-Renom, and L. A. Mirny. Exploring the three-dimensional organization of genomes: interpreting chromatin interaction data. Nature Reviews Genetics, 14(6):390–403, 2013.

[20] M. Imakaev, G. Fudenberg, R. P. McCord, N. Naumova, A. Goloborodko, B. R. Lajoie, J. Dekker, and L. A. Mirny. Iterative correction of Hi-C data reveals hallmarks of chromosome organization. Nature methods, 9(10):999, 2012.

[21] W. Kabsch. A solution for the best rotation to relate two sets of vectors. Acta Crystallographica Section A: Crystal Physics, Diffraction, Theoretical and General Crystallography, 32(5):922–923, 1976.

[22] T. Yang, F. Zhang, G. G. Yardımcı, F. Song, R. C. Hardison, W. S. Noble, F. Yue, and Q. Li. HiCRep: assessing the reproducibility of Hi-C data using a stratum-adjusted correlation coefficient. Genome research, 27(11):1939–1949, 2017.

[23] F. Ay, T. L. Bailey, and W. S. Noble. Statistical confidence estimation for Hi-C data reveals regulatory chromatin contacts. Genome research, 24(6):999–1011, 2014.

[24] M. F. Lyon. Gene action in the X-chromosome of the mouse (Mus musculus L.). nature, 190(4773):372–373, 1961.

[25] T. J. Stevens, D. Lando, S. Basu, L. P. Atkinson, Y. Cao, S. F. Lee, M. Leeb, K. J. Wohlfahrt, W. Boucher, A. O’Shaughnessy-Kirwan, J. Cramard, A. J. Faure, M. Ralser, E. Blanco, L. Morey, M. Sans’, M. G. S. Palayret, B. Lehner, L. D. Croce, A. Wutz, B. Hendrich, D. Klenerman, and E. D. Laue. 3D structures of individual mammalian genomes studied by single-cell Hi-C. Nature, 544(7648):59–64, 2017.

[26] F. Ramírez, V. Bhardwaj, L. Arrigoni, K. C. Lam, B. A. Grüning, J. Villaveces, B. Habermann, A. Akhtar, and T. Manke. High-resolution TADs reveal DNA sequences underlying genome organization in flies. Nature communications, 9(1):1–15, 2018.

[27] H. Yoo-Warren, V. Pachnis, R. S. Ingram, and S. M. Tilghman. Two regulatory domains flank the mouse H19 gene. Molecular and Cellular Biology, 8(11):4707–4715, 1988.

[28] P. A. Leighton, J. R. Saam, R. S. Ingram, C. L. Stewart, and S. M. Tilghman. An enhancer deletion affects both H19 and Igf2 expression. Genes & development, 9(17):2079–2089, 1995.

[29] K. Ishihara, N. Hatano, H. Furuumi, R. Kato, T. Iwaki, K. Miura, Y. Jinno, and H. Sasaki. Comparative genomic sequencing identifies novel tissue-specific enhancers and sequence elements for methylation-sensitive factors implicated in Igf2/H19 imprinting. Genome research, 10(5):664–671, 2000.

[30] J. L. Thorvaldsen, K. L. Duran, and M. S. Bartolomei. Deletion of the H19 differentially methylated domain results in loss of imprinted expression of H19 and Igf2. Genes & development, 12(23):3693–3702, 1998.

[31] D. P. Barlow and M. S. Bartolomei. Genomic imprinting in mammals. Cold Spring Harbor perspectives in biology, 6(2):a018382, 2014.

[32] M. Merkenschlager and D. T. Odom. CTCF and cohesin: linking gene regulatory elements with their targets. Cell, 152(6):1285–1297, 2013.

[33] D. Lleres, B. Moindrot, R. Pathak, V. Piras, M. Matelot, B. Pignard, A. Marchand, M. Poncelet, A. Perrin, V. Tellier, R. Feil, and D. Noordermeer. CTCF modulates allele-specific sub-TAD organization and imprinted gene activity at the mouse Dlk1-Dio3 and Igf2-H19 domains. Genome Biology, 20(1):1–17, 2019.

[34] A. Murrell, S. Heeson, and W. Reik. Interaction between differentially methylated regions partitions the imprinted genes Igf2 and H19 into parent-specific chromatin loops. Nature genetics, 36(8):889–893, 2004.

[35] F. Court, M. Baniol, H. Hagege, J. S. Petit, M. N. Lelay-Taha, F. Carbonell, M. Weber, G. Cathala, and T. Forne. Long-range chromatin interactions at the mouse Igf2/H19 locus reveal a novel paternally expressed long non-coding RNA. Nucleic acids research, 39(14):5893–5906, 2011.

[36] S. Kurukuti, V. K. Tiwari, G. Tavoosidana, E. Pugacheva, A. Murrell, Z. Zhao, V. Lobanenkov, W. Reik, and R. Ohlsson. CTCF binding at the H19 imprinting control region mediates maternally inherited higher-order chromatin conformation to restrict enhancer access to Igf2. Proceedings of the national academy of sciences, 103(28):10684–10689, 2006.

[37] A. C. Bell and G. Felsenfeld. Methylation of a CTCF-dependent boundary controls imprinted expression of the Igf2 gene. Nature, 405(6785):482–485, 2000.

[38] A. T. Hark, C. J. Schoenherr, D. J. Katz, R. S. Ingram, J. M. Levorse, and S. M. Tilghman. CTCF mediates methylation-sensitive enhancer-blocking activity at the H19/Igf2 locus. Nature, 405(6785):486–489, 2000.

[39] J. E. Phillips-Cremins, M. E. G. Sauria, A. Sanyal, T. I. Gerasimova, B.R. Lajoie, J. S. K. Bell, C. T. Ong, T. A. Hookway, C. Guo, Y. Sun, M. J. Bland, W. Wagstaff, S. Dalton, T. C. McDevitt, R. Sen, J. Dekker, J. Taylor, and V. G. Corces. Architectural protein subclasses shape 3D organization of genomes during lineage commitment. Cell, 153(6):1281–1295, 2013.

[40] C. Cubeñas-Potts and V. G. Corces. Topologically associating domains: an invariant framework or a dynamic scaffold? Nucleus, 6(6):430–434, 2015.

[41] E. Crane, Q. Bian, R. P. McCord, B. R. Lajoie, B. S. Wheeler, E. J. Ralston, S. Uzawa, J. Dekker, and B. J. Meyer. Condensin-driven remodelling of X chromosome topology during dosage compensation. Nature, 523(7559):240–244, 2015.

[42] K. Kruse, C. B. Hug, B. Hernández-Rodríguez, and J. M. Vaquerizas. TADtool: visual parameter identification for TAD-calling algorithms. Bioinformatics, 32(20):3190–3192, 2016.

